# A nutrient-sensitive enterokine coordinates developmental plasticity through inter-organ signalling

**DOI:** 10.1101/2025.11.06.687047

**Authors:** L Bai, J Montagne, CI Ramos, F Leulier

## Abstract

Animal survival in fluctuating environments depends on the ability to modulate their developmental pace in response to nutrient availability, a phenomenon known as developmental plasticity. In *Drosophila* larvae, we uncover a critical endocrine mechanism that coordinates this process under conditions of amino acid restriction. We identify the peptide hormone Limostatin as an enterokine, produced by a small population of larval midgut enteroendocrine cells, that acts systemically to inhibit the expression and release of dIlp2, a major insulin-like peptide controlling developmental progression. Limostatin expression and secretion by enteroendocrine cells is triggered by reduced amino acid availability through an inter-organ relay involving the fat body and neuroendocrine insulin-producing cells in the brain. In turn, Limostatin participates in a feedback control loop that slows down developmental progression once systemic nutrient shortage is sensed. This bidirectional gut-brain axis enables larvae to preserve viability under nutritional stress. Our findings define the larval gut as a nutrient-sensitive endocrine organ and position Limostatin as a key regulator of developmental plasticity. Our work expands the concept of decretins to include developmental pace control, suggesting that enterokines that regulate IGF signalling, rather than insulin release per se, may represent an evolutionarily conserved or convergent strategy in regulating developmental plasticity.

## Introduction

Animals must continuously adjust their physiology to survive in environments where nutrient availability fluctuates, and a central aspect of this adaptation is the modulation of developmental pace in response to nutritional input, a process known as developmental plasticity^1^. Among dietary components, protein availability is a major determinant of somatic growth and maturation rate^2^, and protein restriction typically triggers a reduction in growth rate and delayed maturation, an energy-conserving response that helps preserving developmental homeostasis to ensure survival to the adulthood minimizing costs on reproductive capacity^3,4^. In parallel, the intestinal microbiome plays a key role in digestion, nutrient assimilation and endocrine maturation, and is itself sensitive to dietary deprivation^5,6^. Alterations in microbial composition can either buffer or amplify the impact of poor nutrition by modulating host metabolism and hormonal signalling^7,8^. Thus, developmental plasticity is best understood as a multilayered physiological response, integrating environmental cues such as diet and microbiota with internal states like metabolism and endocrine regulation to coordinate developmental timing and survival^9^.

*Drosophila melanogaster* offers a genetically tractable model to dissect how these signals coordinate developmental pace with nutrient availability. Larval development in *Drosophila* is highly plastic, tightly regulated by environmental conditions, and largely orchestrated by insulin-like peptides (dIlps), which function as the principal somatotropic signals in insects^10,11^. dIlps can be produced locally^12^ by a cluster of Insulin-producing cells (IPCs) in the brain, which act systemically through the insulin receptor (InR) in peripheral tissues^13^. One key downstream target of InR signalling is the transcription factor FoxO, which represses anabolic gene programs when dIlp signalling is reduced^14,15^. The output of IPCs’ activity is tightly regulated by a network of inter-organ communications^16^. Nutrient uptake by enterocytes in the gut, amino acid sensing via TORC1 in the fat body - a tissue that fulfils hepatic and adipose functions^17^ - and fat body-derived adipokine signals^18^ including Upd2^19^, Stunted^20^, Eiger^21^ and GBP1/2^22^ fine-tune dIlp production by IPCs to match nutritional status. Alongside these systemic cues, the intestinal microbiome contributes to developmental pace by producing limiting nutrients^23^, facilitating digestion^24^, and influencing endocrine pathways, including dIlps signalling^25,26^. Together, these findings highlight the complex cross-talk between diet, microbiota, and endocrine organs in shaping developmental growth and maturation.

Previous work identified the peptide hormone Limostatin (Lst) as a fasting-induced “decretin” hormone that suppresses dIlp production and release in adult flies^27^. Lst is produced by the corpora cardiaca, an endocrine tissue producing adipokinetic hormone (AKH)–the insect analog of glucagon–and its upregulation under sugar deprivation contributes to reduced systemic insulin signalling and increased glycemia. Although overexpression of *Lst* in the larval fat body was reported to delay development, *Lst* loss-of-function animals do not display overt developmental timing phenotype under standard laboratory conditions^27^, raising questions about its developmental role and tissue-specific function. Here, we identify a novel, developmentally relevant role for Lst as a gut-derived hormone that controls developmental pace and survival in response to nutritional challenges. We show that Lst is specifically induced in a discrete group of enteroendocrine cells (EECs) in the anterior midgut of larvae exposed to extreme amino acid stress or milder amino acid stress coupled with microbiota depletion. We show that Lst production and release by EECs is induced upon reduced circulating dIlp2 levels, and that Lst in turn suppresses dIlp2 release from brain IPCs. This establishes an amplifying regulatory loop that reinforces the adaptation of developmental pace to amino acid scarcity. These findings establish that in contrast to its role as a decretin regulating sugar homeostasis in adults, Lst in juveniles acts as an enterokine, a gut-derived peptide hormone that integrates amino acid signals of dietary and microbial origins to coordinate developmental plasticity.

## Results

### Lst is a peptide hormone regulating developmental progression and survival under malnutrition

In our recent study^28^, we performed transcriptomic profiling of dissected larval midguts from *Drosophila* reared on a low-yeast diet, a model of malnutrition that elicits developmental adaptation, characterized by reduced larval growth rate and delayed pupariation in wild-type (WT) animals. Given the known role of the microbiome in buffering nutritional stress, we compared germ-free larvae with animals mono-associated with *Lactiplantibacillus plantarum^WJL^* (*Lp^WJL^,* referred as *Lp* here), a well-characterized member of the *Drosophila* gut microbiota that recapitulates the growth-promoting effects of a complex microbiome under malnutrition^25^. Among the differentially regulated transcripts, we identified *Limostatin* (*Lst*) as significantly upregulated in germ-free larvae raised under malnutrition (Fig. 1A)^28^. Lst was previously characterized in adult flies as a peptide hormone secreted by corpora cardiaca during fasting, where it suppresses dIlp production and release from the brain^27^. Our data suggest that Lst may also contribute to the adaptive growth response of germ-free larvae exposed to malnutrition. To test this hypothesis, we analyzed *Lst* loss-of-function mutants. In line with previous findings, heterozygous and homozygous *Lst* null mutant (*Lst*^1^) animals showed normal developmental progression under standard nutritional conditions^27^ (Fig. 1B) but a severe developmental lethality was observed in homozygous mutants (Fig. 1C).

**Figure 1.**
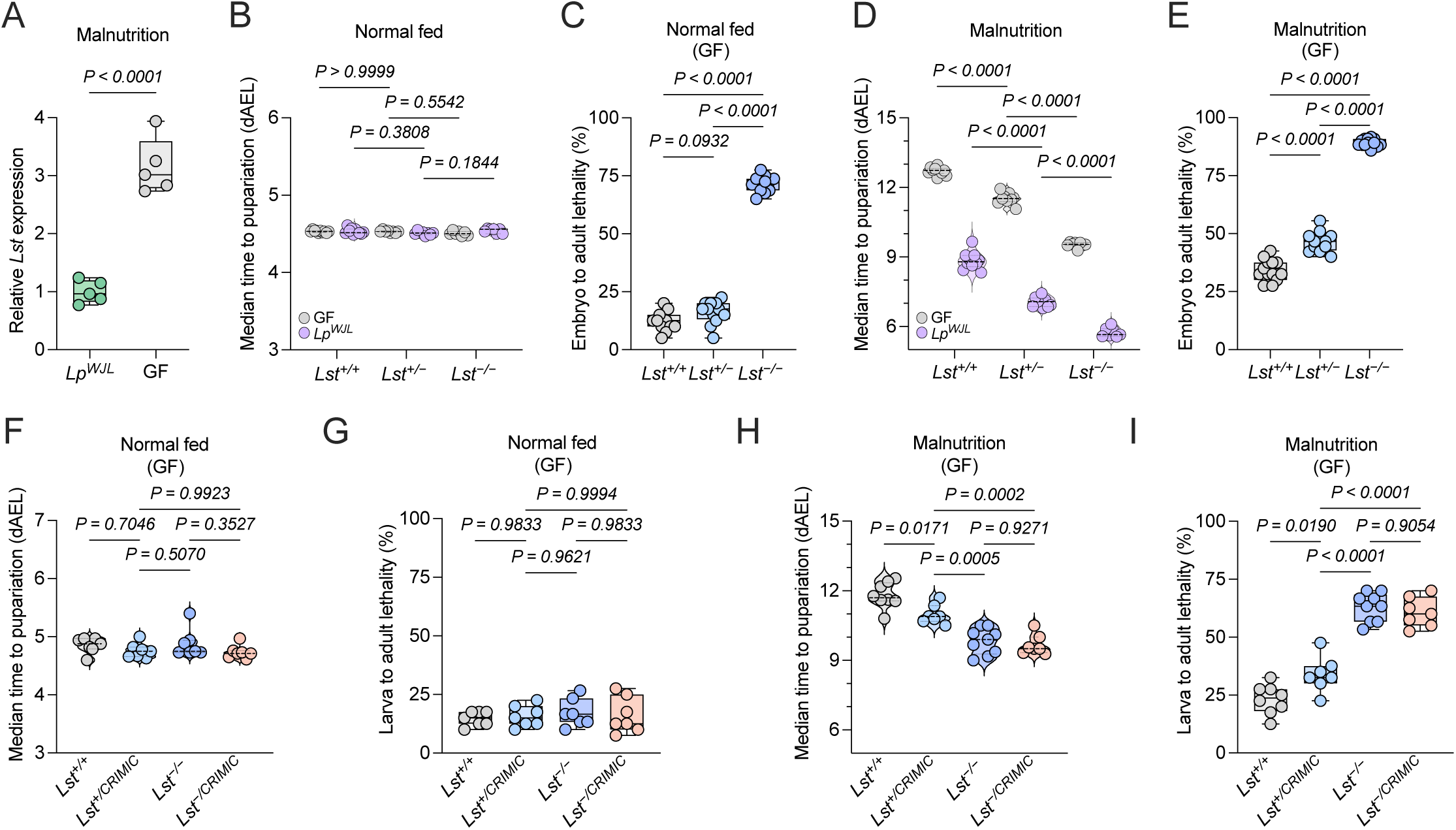
Lst is a peptide hormone regulating adaptive growth and survival under malnutrition. (A) Transcript levels of gut-derived *Lst* analyzed via midgut bulk RNA-seq data from germ-free (GF) and *Lactiplantibacillus plantarum* (*Lp*)-associated undernourished *yw* larvae (*n* = 5). The complete RNA-seq dataset and statistical analyses are available in Bai et al. 2026 ^28^ and deposited in the NCBI Gene Expression Omnibus (GEO) database under accession number GSE310159. (B–E) Median time to pupariation (denoted as days after egg laying, dAEL) (B, D) and percentage of embryo-to-adult lethality (C, E) measured for homozygous and heterozygous *Lst* null mutants, alongside control larvae. Assessments were performed under normal fed conditions (B, C) or under a 7 g/L low-yeast malnutrition diet (D, E) (*n* ≥ 7). (F–I) Median time to pupariation (F, H) and percentage of L1-to-adult lethality (G, I) measured across genotypes, including trans-heterozygous *Lst* mutants, heterozygous mutants carrying a *CRIMIC* insertion, homozygous *Lst* null mutants, and control larvae. Assessments were performed under normal fed conditions (F, G) or a 7 g/L low-yeast malnutrition diet (H, I) (*n* ≥ 7). Statistical significance was determined using a two-tailed unpaired *t* test (A), a two-way ANOVA with Tukey’s multiple comparisons test (B, D), or a one-way ANOVA with Tukey’s multiple comparisons test (C, E, F–I). Exact *P* values are indicated on the respective panels.

However, under malnourished conditions (low yeast diet, 7 g/L), both heterozygous and homozygous *Lst* null mutants exhibited significantly accelerated development (Fig. 1D). Yet, heterozygous animals displayed reduced viability during development and homozygous animals had high lethality (Fig. 1E) relative to their WT siblings and normal nutrition controls.

To determine whether Lst is specifically required for viability under malnutrition, we adopted a trans-heterozygous strategy by crossing the *Lst*^1^ null allele (*Lst*^−/−^) with a second allele carrying a *CRIMIC* insertion within the *Lst* locus, located at the beginning of the sole intron of the *Lst* gene (*Lst^CR018^*^53^, hereafter referred to as *Lst^CRIMIC^*). To bypass potential early embryonic lethality, we specifically selected first-instar (L1) larvae of each genotype and monitored larval-to-adult development and viability. Strikingly, upon normal nutrition, both *Lst*^−/−^ and *Lst^−/CRIMIC^* larvae displayed normal developmental timing (Fig. 1F) and survival rates (Fig. 1G) comparable to controls once they enter the L1 stage after hatching. These results indicate that the reduced viability initially observed upon normal nutrition in *Lst^−/−^* animals (Fig. 1C) results from an embryonic lethality and/or reduced fertilization/oogenesis efficiency in this specific homozygous context. Importantly, under malnutrition conditions, all mutant genotypes (*Lst^CRIMIC/+^* - *Lst^CRIMIC/^*^−^ - *Lst*^−*/*−^) exhibited significantly accelerated developmental timing (Fig. 1H) coupled with an increased lethality (Fig. 1I). These findings strengthen our conclusion that Lst is a critical regulator of adaptive larval growth, restraining developmental pace in order to preserve larval viability during nutrient stress.

These phenotypes were independently confirmed using two distinct *Lst*-targeting RNAi lines driven by *Lst-GAL4* driver^27^ (Fig. S1A–D). Importantly, despite their accelerated developmental progression, *Lst*-RNAi animals did not exhibit increased food intake (Fig. S1E–F), indicating that the enhanced developmental rate is not attributable to elevated feeding activity. Together, these findings establish Lst as a peptide hormone that coordinates developmental progression and survival in response to nutritional stress and microbiome depletion. Its expression under these conditions suggests a role as a critical effector in the endocrine network governing adaptive developmental timing.

### Nutritional stress activates Lst in anterior midgut EECs

Prompted by the upregulation of *Lst* transcripts in dissected midguts from germ-free larvae raised under malnutrition (Fig. 1A), we examined the tissue-specific expression of *Lst* using a *Lst-GAL4*>GFP reporter line^27^. As previously reported in adults, *Lst-GAL4* drives GFP expression in the larval corpora cardiaca (Fig. 2A, Fig. S2A). However, we also detected additional GFP-positive cells in a discrete region of the anterior midgut, just anterior to the copper cell region (Fig. 2B–C, Fig. S2B), a signal we did not detect in adult midguts. Although we cannot exclude the possibility that the *Lst-GAL4* driver does not fully recapitulate endogenous *Lst* expression, all GFP-positive cells in this region co-localized with Prospero, a marker of EECs (Fig. 2B-C). Of note, GFP-labelled cells were basally located in the epithelium with no obvious apical projections to the intestinal lumen (Movie S1). We observed less than 10 GFP-positive cells per midgut with no marked differences across diet or microbiome conditions (Fig. 2D), indicating a restricted and stereotyped expression pattern. To further confirm endogenous *Lst* expression, we generated affinity-purified antibodies targeting a 15-amino acid peptide of the Lst protein. Immunostaining with these antibodies revealed specific signal in the same anterior EEC subset; notably, this signal was completely abolished in the midguts of *Lst^1^* animals, hereafter referred to as the Lst⁺ EEC population (Fig. 2B–C, Fig. S3A).

**Figure 2.**
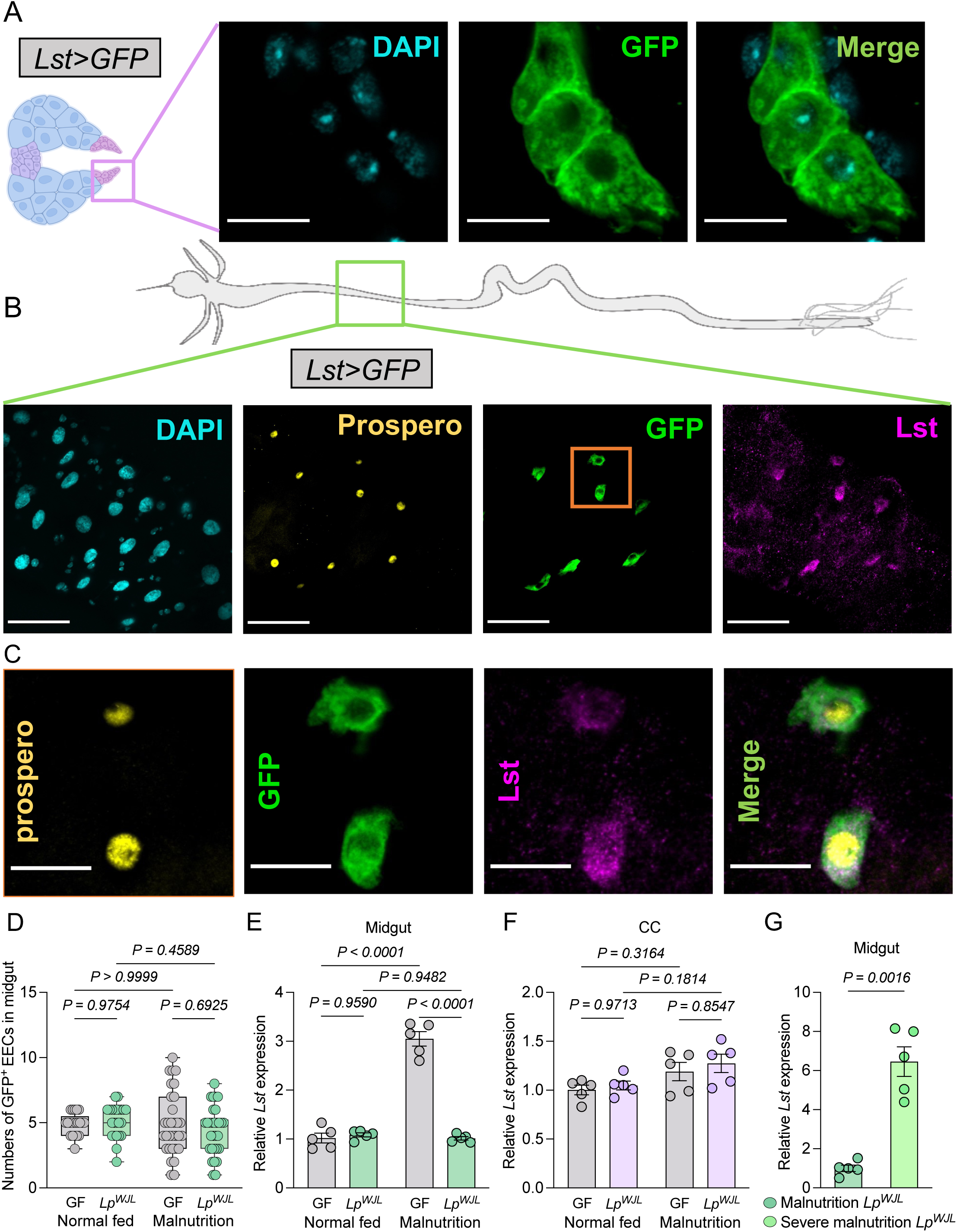
Lst is induced in a discrete population of anterior midgut enteroendocrine cells under malnutrition and microbiota depletion (A, B) *Lst* expression revealed by the *Lst-GAL4>GFP* reporter line under germ free (GF) and malnourished conditions. The reporter line reveals *Lst* expression in the corpora cardiaca (CC) of larvae (A), consistent with adult expression patterns. (B) GFP-positive cells co-localize with the EEC marker Prospero in the midgut region anterior to the acidic zone. Immunostaining confirms the presence of gut-derived Lst in larvae. (C) Magnified views of the selected orange region from panel B highlighting the co-localization of Prospero, Lst peptide and anti-GFP staining within these EECs. (D) Quantification of Lst-positive EECs in the larval midgut using the *Lst-GAL4>GFP* reporter line (*n* ≥ 21). (E–G) Transcript levels of gut-derived *Lst* (E, G) and corpora cardiaca-derived *Lst* (F) analyzed by qRT-PCR (*n* = 5). Malnutrition and severe malnutrition correspond to 7 g/L and 3 g/L yeast diets, respectively. Scale bars: 5 µm in (A), 50 µm in (B), and 15 µm in (C). In (E–G), data are presented as mean ± SEM. Statistical significance was determined using a two-way ANOVA with Tukey’s multiple comparisons test (D–F) or a two-tailed Welch’s *t* test (G). Exact *P* values are indicated on the respective panels.

To investigate the regulation of *Lst* expression, we performed RT-qPCR analyses on dissected midguts and corpora cardica from larvae raised under either conventional or malnutrition diets, in both germ-free and *Lp*-associated conditions (Fig. 2E–F). These experiments revealed a robust and selective induction of *Lst* expression in the midgut of malnourished germ-free animals, while in contrast to adults^27^, expression in the corpora cardiaca remained unchanged. We further found that severe dietary restriction alone was sufficient to trigger strong induction of *Lst* expression, even in the absence of microbiota alteration: *Lst* transcript levels were increased approximately six-fold in the midguts of *Lp***-**associated larvae raised on an extremely low-yeast diet (3 g/L) compared to those on a low-yeast diet (7 g/L) (Fig. 2G).

To extend these findings at the protein level, we next performed complementary immunostaining analyses using anti-Lst antibody. We detected Lst protein in the relevant EEC population, and observed differences in steady-state intracellular staining across nutritional and microbiome conditions that paralleled the transcriptional changes (Fig. 3A). We then sought to visualize intracellular accumulation of Lst by acutely blocking vesicle-mediated secretion. To this end, we expressed the temperature-sensitive dynamin allele *shibire^ts^* ^29^ in Lst^+^ cells and shifted larvae to the restrictive temperature for 3 hours to inhibit secretion. Under these conditions, we observed a marked intracellular accumulation of Lst protein, which was substantially stronger in undernourished germ-free larvae than in normally fed controls (Fig. 3B). Together, these experiments support the conclusion that the induction of *Lst* mRNA under malnutrition is accompanied by increased Lst protein production and secretion. Together, these results identify a previously unrecognized site of Lst expression during larval stage and demonstrate that Lst is specifically expressed and secreted by this discrete population of anterior midgut EECs in response to combined mild nutritional stress and microbiota depletion or to severe malnutrition alone.

**Figure 3.**
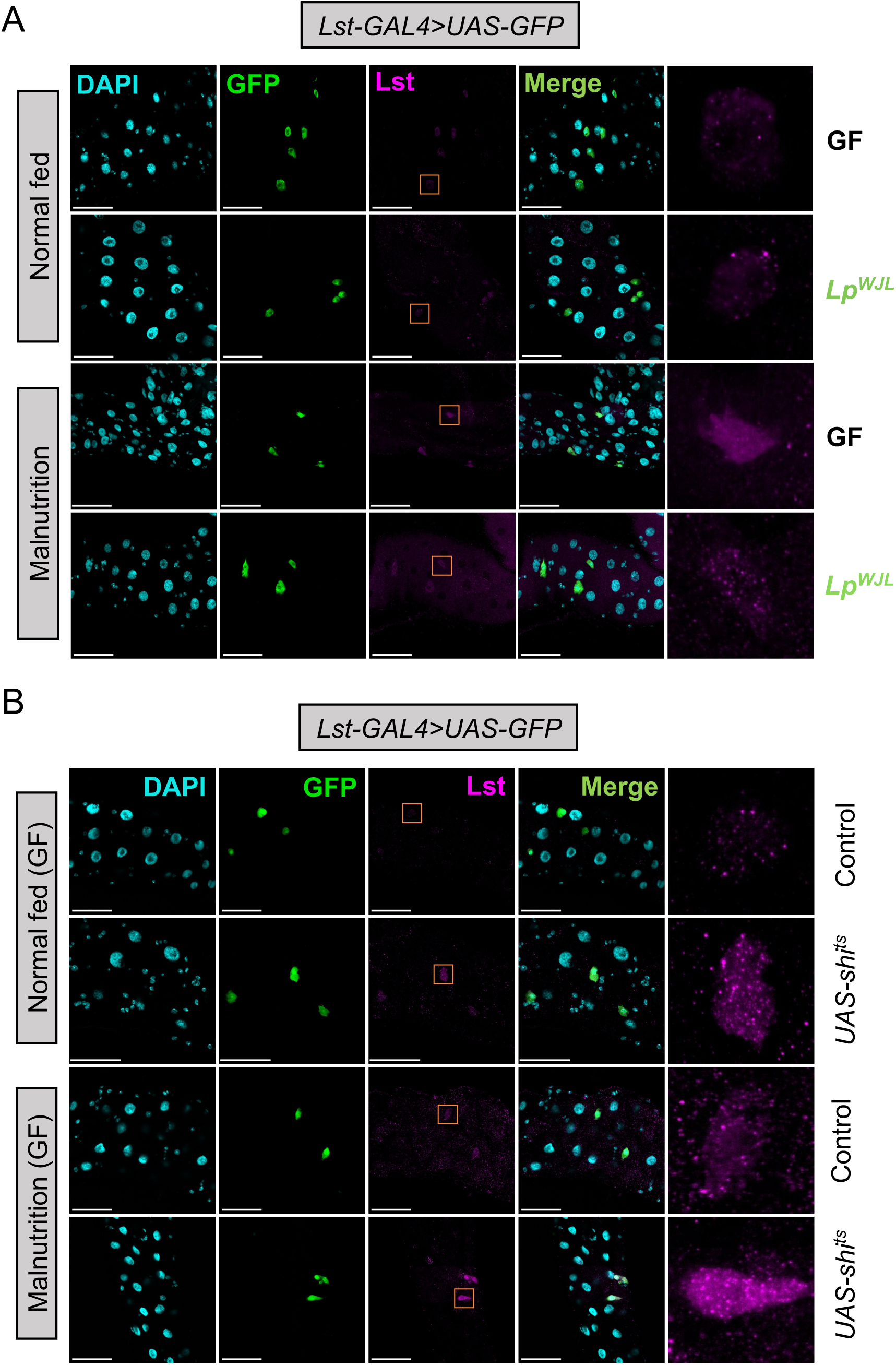
Lst peptide adaptive levels in the larval midgut (A) Lst immunostaining in the *Lst-GAL4>GFP* reporter line of germ free and *Lp*-associated larvae under normal fed or malnourished conditions. Magnified views of the selected orange regions highlight endogenous Lst levels. (B) Lst immunostaining in germ free control (*Lst-GAL4>GFP*) or experimental (*Lst-GAL4>GFP; UAS-shi^ts^)* larvae under normal fed or malnourished conditions. Mid-third-instar (L3) larvae were shifted from 25°C to 29°C for 3 hours to restrict Lst peptide secretion. Magnified views of the selected orange region highlight intracellular Lst accumulation. Scale bars: 50 µm in (A, B).

### Lst from EECs regulates adaptive growth and survival under malnutrition and microbiota depletion

We next sought to determine which pool of Lst - produced by the corpora cardiaca and/or by EECs - contributes to the regulation of adaptive growth under nutritional and microbiome stress. Consistent with earlier findings showing that ectopic *Lst* expression in the larval fat body can delay developmental progression^27^, we found that targeted overexpression of *Lst* in either EECs (using an EEC-specific GAL4 driver; *EEC*-GAL4^30^; Fig. 4A) or corpora cardiaca cells (using the *AKH*-GAL4 driver^31^; Fig. 4B) was sufficient to impair developmental progression in larvae mono-associated with *Lp*. These results indicate that Lst is sufficient to modulate maturation rates, independently of its tissue of origin.

**Figure 4.**
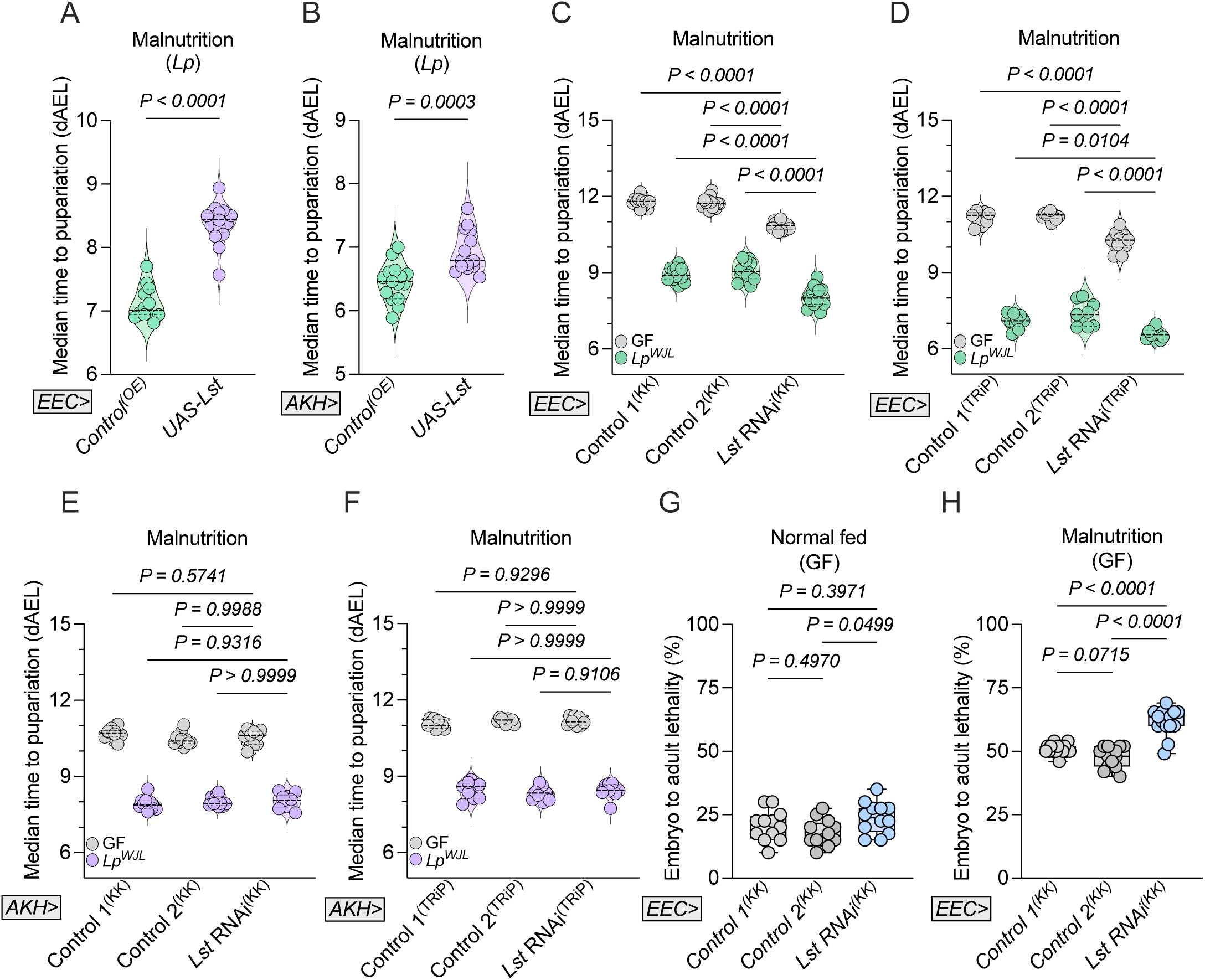
Gut-derived Lst regulates adaptive growth and survival under malnutrition and microbiota depletion (A–F) Median time to pupariation measured under malnutrition (7 g/L low-yeast diet) for *Lp*-associated larvae overexpressing *Lst* in EECs (*EEC>UAS-Lst*) (A) or in corpora cardiaca (CC) (*AKH>UAS-Lst*) (B) with their respective overexpression controls, as well as *Lst* knockdown in EECs (*EEC>UAS-Lst* RNAi) or in corpora cardiaca (*AKH>UAS-Lst* RNAi), using either the KK RNAi line (C, E) or the TRiP RNAi line (D, F) compared to their respective KK or TRiP controls (*n* ≥ 8). (G, H) Embryo-to-adult lethality was quantified for EEC-specific *Lst* knockdown larvae (*EEC>Lst* RNAi) and their KK controls under normal fed conditions (G) or under malnutrition (H) (*n* ≥ 12). Statistical significance was determined using a two-tailed unpaired *t* test (A, B), two-way ANOVA with Tukey’s multiple comparisons test (C–F), or a one-way ANOVA with Tukey’s multiple comparisons test (G, H). Exact *P* values are indicated on the respective panels.

To test which Lst source is necessary for adaptive growth regulation under nutritional stress, we used tissue-specific RNA interference to knock down *Lst* expression. We expressed two independent RNAi constructs targeting *Lst* in either EECs (*EEC-GAL4*, Fig. 4C–D, Fig. S3B–C, Fig. S4A–B or *Voilà-GAL4*^32^, Fig. S4C) or in corpora cardiaca cells (*AKH-GAL4*, Fig. 4E–F, Fig. S4D–E). These experiments were performed in both germ-free and *Lp*-associated larvae, under either conventional or malnutrition diets (7 g/L low-yeast diet). Strikingly, *Lst* knockdown in EECs (Fig. 4C–D, G–H, Fig. S4A–C), but not in corpora cardiaca cells (Fig. 4E–F, Fig. S4D–E) was sufficient to phenocopy the altered adaptive maturation (Fig. 4C–D, Fig. S4A–C) and low viability (Fig. 4G–H) observed in whole-animal *Lst* null heterozygous and trans-heterozygous mutants (Fig. 1C–E, H, I) under germ-free malnutrition conditions.

Importantly, despite their accelerated developmental progression, EEC>*Lst*-RNAi animals did not exhibit increased food intake (Fig. S4F–G). These findings demonstrate that Lst derived from EECs is necessary for restraining developmental progression and ensuring viability when animals are challenged by both nutrient deprivation and microbiota loss.

### Lst expression in EECs is sensitive to amino acid of dietary and microbial origins

Our findings indicate that Lst expression in EECs is modulated by fluctuations in both diet and microbiota. While Lst was previously shown to be regulated by dietary sugar levels in adult corpora cardiaca cells^27^, it is unclear which nutrients influence its expression in the larval midgut. Notably, conventional inactivated yeast-based (oligidic) diets do not allow for a precise dissection of the nutritional components responsible for Lst regulation, as yeast provides a complex mixture of macro-and micronutrients. Although commonly referred to as “low-protein” diets, yeast-reduced regimens likely affect multiple nutritional inputs beyond amino acids availability. To directly test whether dietary amino acids regulate Lst expression and function, we employed the FLYAA holidic diet system, a chemically defined nutritional framework previously established for dissecting nutrient-microbiota-host interactions in *Drosophila*^23,33^. Within this system, we altered total amino acid content while maintaining their relative proportions. Specifically, we used a High Amino Acid Holidic Diet (HAAHD; 21.38 g/L total AAs) and a Low Amino Acid Holidic Diet (LAAHD; 8.55 g/L total AAs), the latter of which has been shown to impair developmental growth^23^. Using these holidic diets, we found elevated *Lst* transcripts levels in the midgut (Fig. 5A) but not in corpora cardiaca (Fig. S5A) under amino acid restriction and microbiota depletion, mirroring our previous observations in germ-free animals fed oligidic low-yeast diets. Functionally, we confirmed that Lst is required for coordinating development under these conditions: germ-free *Lst¹* mutants (Fig. 5B) and germ-free animals subjected to tissue-specific *Lst* knockdown in EECs (Fig. 5C–D, Fig. S5B–C), but not in corpora cardiaca cells (Fig. S5D–G), exhibited altered adaptive maturation under the LAAHD regimen. Together, these findings demonstrate that Lst acts as a key hormonal regulator of adaptive developmental growth in response to dietary amino acid availability and microbiota status.

**Figure 5.**
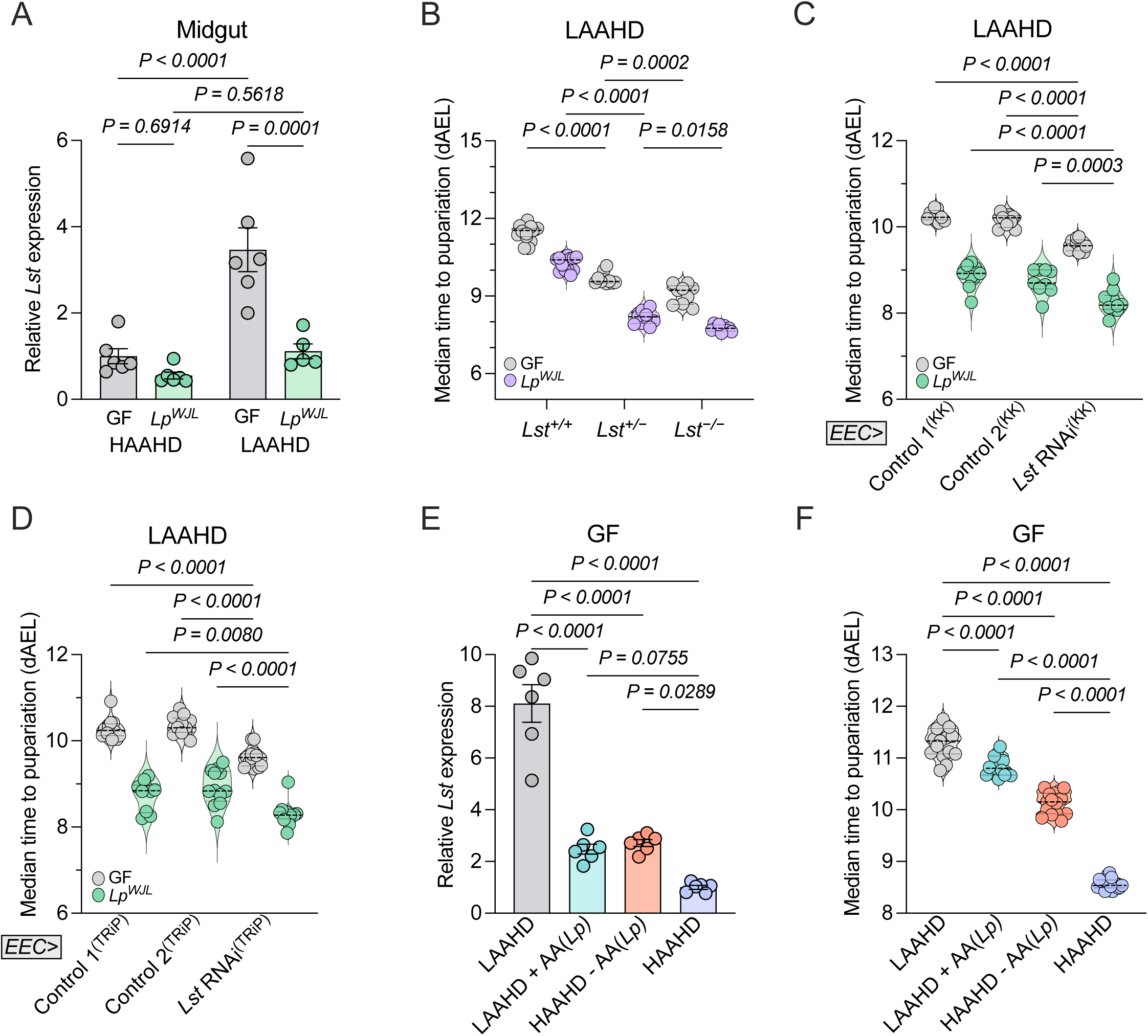
Lst expression in EECs is sensitive to dietary and microbiota-derived AAs (A) Relative transcript levels of gut-derived *Lst* analyzed by qRT-PCR in germ-free (GF) and *Lp*-associated larvae (*Lp*) reared under HAAHD and LAAHD (*n* ≥ 5). Data are presented as mean ± SEM. (B) Median time to pupariation measured for homozygous and heterozygous *Lst* null mutants, alongside control larvae reared under LAAHD (*n* ≥ 9). (C, D) Median time to pupariation measured under LAAHD for larvae with *Lst* knockdown in EECs (EEC>UAS-*Lst* RNAi) using the *KK* RNAi line (C) and the *TRiP* RNAi line (D), each compared against its respective controls (*n* ≥ 9). (E, F) Relative transcript levels of gut-derived *Lst* analyzed by qRT-PCR (E) and median time to pupariation (F) of larvae reared in HDs where specific AAs produced by *Lp* were either supplemented to LAAHD or reduced from HAAHD. *n* = 6 (E), *n* ≥ 15 (F). Statistical significance was determined using either a two-way ANOVA (A–D) or a one-way ANOVA (E, F) followed by Tukey’s multiple comparisons test. Exact *P* values are indicated on the respective panels.

We next explored whether microbiota-mediated nutrient provisioning contributes to Lst repression. Given the role of amino acid availability in regulating Lst expression, and our prior demonstration that commensal bacteria, including *Acetobacter pomorum (Ap)* and *Lp* strains, can supplement amino acids to their juvenile host during malnutrition^23^, we hypothesized that *Lp* represses Lst expression by supplying limiting amino acid s. To test this hypothesis, we raised germ-free larvae on either HAAHD or LAAHD that were selectively depleted or supplemented with the specific amino acids that *Lp* is able to produce (His, Lys, Met, Phe, Thr and Asp. Fig. 5E–F). Consistent with our model, *Lst* expression was significantly repressed and developmental progression improved when His, Lys, Met, Phe, Thr and Asp were added to the LAAHD (Fig. 5E–F). Conversely, reducing the concentration of those same AAs from the HAAHD to their levels in LAAHD led to Lst induction and maturation delay (Fig. 5E–F). Similarly, association of germ-free larvae raised on LAAHD with *Ap,* a commensal bacteria capable of producing all amino acids, antagonized *Lst* expression upon malnutrition (Fig. S5H), indicating that microbial production of amino acids may contribute to Lst-dependent developmental growth regulation.

### Cellular amino acid sensors are dispensable for Lst induction in EECs

We next sought to identify how amino acid restriction leads to Lst induction specifically in EECs. A straightforward hypothesis is that Lst⁺ EECs directly sense amino acid scarcity in the gut lumen through conserved intracellular nutrient-sensing pathways. Two major evolutionarily conserved signalling hubs govern cellular responses to amino acid availability: the Gcn2 kinase, which is activated by uncharged tRNAs and signals amino acid deprivation^34^, and the target of rapamycin complex 1 (TORC1), a central regulator of growth and metabolism activated by intracellular amino acids and negatively regulated by the Tsc1/2 complex^35^. To assess the functional requirement for these pathways in Lst⁺ EECs, we used EEC-specific RNAi to knock-down *Tsc1* or *Gcn2* (Fig. 6A–B, Fig. 6C–D, respectively). In both cases, *Lst* induction (Fig. 6A,C) and developmental maturation rates (Fig. 6B,D) in undernourished germ-free larvae remained unaffected, indicating that neither Gcn2 nor repression of TORC1 signalling in EECs contribute to *Lst* expression under amino acid-restricted conditions. Together, these results suggest that *Lst* induction in response to amino acid limitation occurs independently of the canonical intracellular amino acid sensors Gcn2 and TORC1 in EECs.

**Figure 6.**
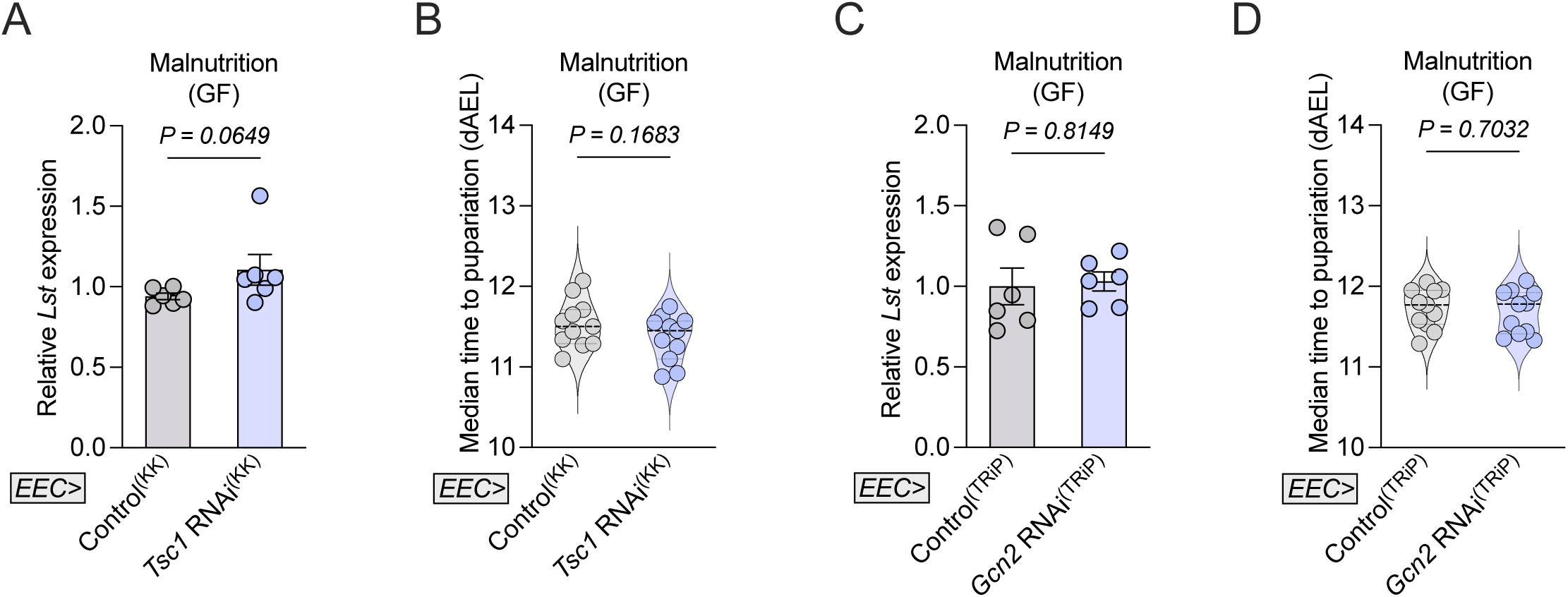
Cellular amino acid sensors are dispensable for Lst induction in EECs (A–D) Under malnutrition, relative transcript levels of gut-derived *Lst* were analyzed by qRT-PCR (A, C) and median time to pupariation was measured (B, D) in germ free (GF) larvae with EEC-specific *Tsc1* knockdown (A, B) or *Gcn2* knockdown (C, D). *n* = 6 for (A, C), *n* = 11 for (B, D). In (A, C), data are shown as mean ± SEM. Statistical significance was determined using a two-tailed Mann-Whitney *U* test (A), two-tailed unpaired *t* test (B–D). Exact *P* values are indicated on the respective panels.

### Lst induction in EECs is controlled by reduced systemic insulin signalling via FoxO

Since *Lst* expression in EECs is not regulated by classical intracellular amino acid sensors such as Gcn2 or TORC1, we next explored whether nutrient-sensitive systemic endocrine signals might be involved. In *Drosophila*, the insulin/insulin-like growth factor signalling (IIS) pathway is a central hormonal regulator of nutrient-dependent growth, and its activity is markedly reduced under protein malnutrition. Decrease in IIS leads to the nuclear translocation of the transcription factor FoxO, which drives transcriptional responses to nutrient stress^36^. To test whether reduced IIS signalling and FoxO activity regulate *Lst* expression, we first monitored intestinal expression of *InR*, a transcriptional target of FoxO and a sensitive negative marker of IIS activity. *InR* transcripts were elevated in dissected midguts of germ-free larvae raised on a low-yeast diet compared to *Lp*-associated controls (Fig. 7A), indicating elevated FoxO activity, hence reduced IIS signalling, in the intestinal epithelium of germ-free animals under malnutrition. Consistently, phosphorylated protein kinase B (p-Akt) immunoreactivity, a marker of IIS pathway activation^37^, was increased in Lst**⁺** EECs of *Lst*>*GFP* larvae, and likely also in some adult midgut precursors, in response to *Lp* association, whereas surrounding enterocytes showed no detectable changes (Fig. 7B).

**Figure 7.**
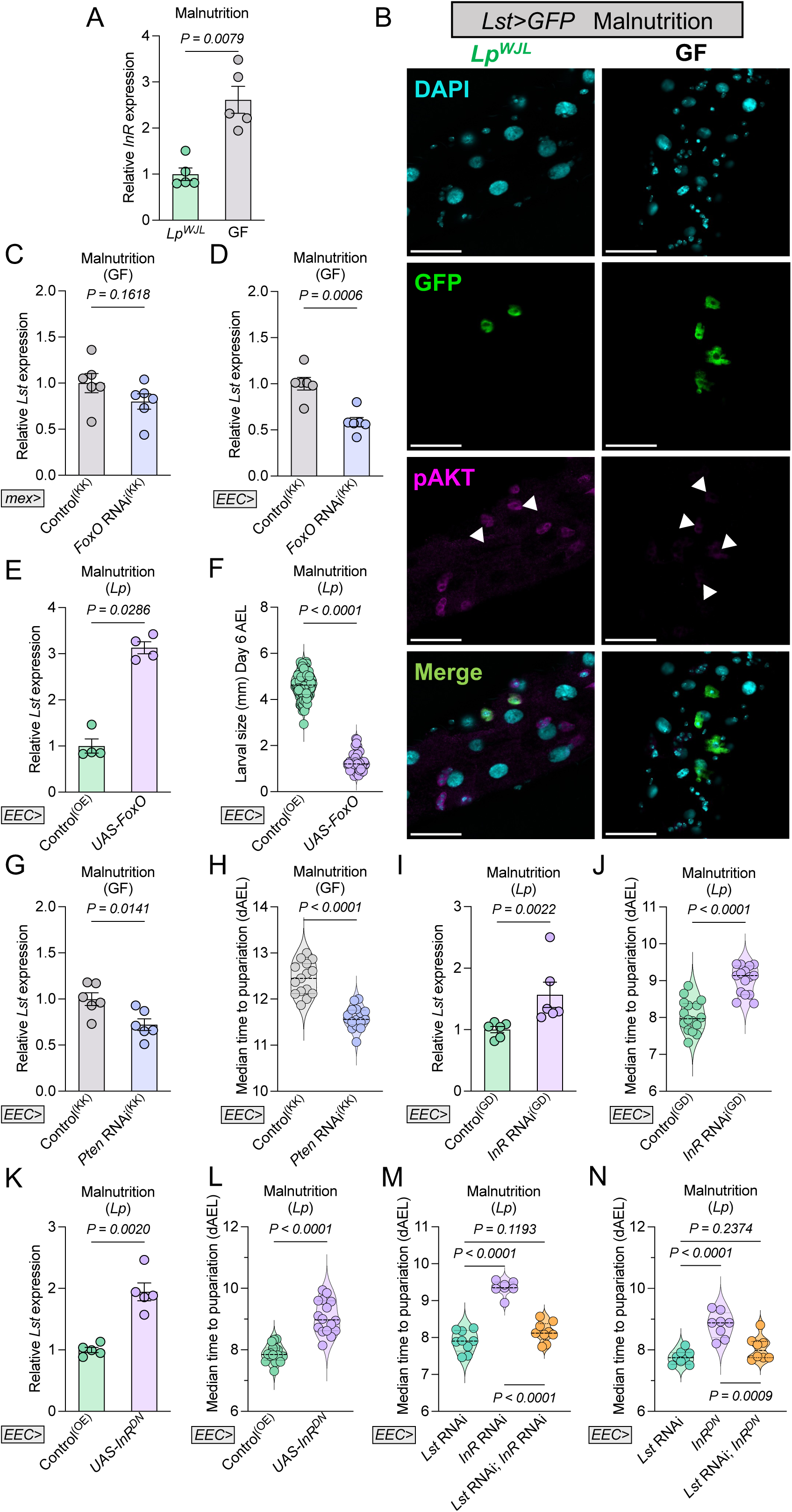
Lst induction in EECs is controlled by reduced systemic insulin signalling via FoxO (A) Relative transcript levels of gut-derived *InR* analyzed by qRT-PCR in *Lp*-associated (*Lp*) and germ free (GF) larvae reared under malnutrition (*n* = 5). (B) Midgut pAKT (phosphorylated AKT) immunostaining. pAKT levels were compared between undernourished germ free and *Lp*-associated *Lst-GAL4>GFP* larvae. Scale bar: 50 μm. (C, D) Under malnutrition, relative transcript levels of gut-derived *Lst* analyzed by qRT-PCR for germ free larvae with *FoxO* knockdown in enterocytes using the *mex-GAL4* (*mex>UAS-FoxO* RNAi) (C), or in EECs using the *EEC-GAL4* (*EEC>UAS-FoxO* RNAi) (D) (*n* = 6). (E, F) Under *Lp*-associated conditions, relative transcript levels of *Lst* analyzed by qRT-PCR (E) and larval sizes measured (F) under malnutrition in larvae overexpressing *FoxO* in EECs (*EEC>UAS-FoxO*) alongside their respective overexpression controls. *n* = 4 (E), *n* ≥ 42 (F). (G, H) Relative transcript levels of gut-derived *Lst* analyzed by qRT-PCR (G) and median time to pupariation measured (H) under malnutrition for germ free larvae with *Pten* knockdown in EECs (*EEC > UAS-Pten* RNAi) and their respective KK controls. *n* = 6 (G), *n* ≥ 15 (H). (I–L) Under malnutrition, relative transcript levels of gut-derived *Lst* analyzed by qRT-PCR (I, K) and median time to pupariation measured (J, L) for *Lp*-associated larvae with *InR* knockdown in EECs (*EEC>UAS-InR* RNAi) and their respective GD controls (I, J), or with EEC-specific overexpression of dominant-negative *InR* (*EEC>UAS-InR^DN^*) and their corresponding overexpression controls (K, L). *n* = 6 (I), *n* = 5 (K), *n* ≥ 15 (J, L). (M, N) Under malnutrition, median time to pupariation measured for germ free larvae with EEC-specific knockdown of both *Lst* and *InR* (*EEC>UAS-Lst* RNAi;*UAS-InR* RNAi) and their respective controls (M), or with *Lst* knockdown combined with overexpression of dominant-negative *InR* (*EEC>UAS-Lst* RNAi; *UAS-InR^DN^*) and their respective controls (N) (*n* ≥ 7). In (A, C–E, G, I, K), data are presented as mean ± SEM. Statistical significance was determined using a two-tailed Mann-Whitney *U* test (A, E, F, I), a two-tailed unpaired *t* test (C, D, G, H, J), a two-tailed Welch’s *t* test (K, L) or a one-way ANOVA followed by Tukey’s multiple comparisons test (M, N). Exact *P* values are indicated on the respective panels.

We next tested whether FoxO activity is required for Lst induction. Tissue-and cell type-specific RNA interference against *FoxO* in enterocytes (*mex*-GAL4; Fig. 7C), EECs (EEC-GAL4; Fig. 7D), and Lst⁺ cells (*Lst*-GAL4; Fig. S6A) revealed that *FoxO* knockdown in EECs or in Lst⁺ cells, but not in enterocytes, reduced *Lst* expression in germ-free animals reared under malnutrition. Conversely, ectopic expression of FoxO in EECs under microbiota-replete conditions was sufficient to drive *Lst* expression (Fig. 7E) and significantly repress larval growth (Fig. 7F). These results establish that FoxO acts cell-autonomously in Lst⁺ EECs to mediate *Lst* induction in response to reduced systemic IIS activity. Furthermore, we analyzed *Lst* genomic locus sequence for potential FoxO response elements using the consensus binding sequences previously described by Spellberg et al.^38^. This analysis identified three putative FoxO-binding motifs within the 2 kb proximal promoter region of *Lst*, as well as an additional consensus motif located within the unique intronic region of the *Lst* gene (Fig. S7). While the presence of these motifs does not formally demonstrate direct binding, these sequence analyses are consistent with a model in which FoxO directly regulates *Lst* transcription in response to nutritional stress.

To further define the regulatory cascade, we manipulated upstream IIS components. RNAi-mediated knockdown of *Pten* in EECs (*EEC-GAL4*; Fig. 7G–H) or Lst⁺ cells (*Lst-GAL4*; Fig. S6B–C) blunted *Lst* induction and accelerated larval development in germ free animals raised on poor diet confirming that enhanced IIS represses *Lst* expression. Conversely, reducing InR activity by expressing *InR* RNAi constructs (Fig. 7I–J; Fig. S6D–E) or a dominant-negative form of InR (*InR^DN^* ^39^; Fig. 7K–L; Fig. S6F–G) in EECs or in Lst⁺ cells robustly induced *Lst* expression and delayed developmental progression (Fig. 7I–L). We next performed genetic epistasis experiments whereby blocking insulin signalling in the EECs—via either *InR*-RNAi or overexpression of a dominant-negative InR (*InR^DN^*)—and *Lst* expression resulted in a growth phenotype that mirrors the loss of *Lst* per se (Fig. 7M–N). These results demonstrate that *Lst* interference is genetically dominant over *InR* loss of function hence indicate that Lst acts downstream of InR in this context. Taken together, these results demonstrate that *Lst* expression in Lst^+^ EECs is negatively controlled by IIS and positively by FoxO, suggesting a direct endocrine link between systemic nutrient sensing and Lst production in the gut.

### Lst induction in EECs is controlled by an inter-organ nutrient sensing regulatory loop

The IIS-dependent control of *Lst* raised the question of how nutrient availability modulated IIS in Lst^+^ EECs. In *Drosophila*, the fat body serves as a central nutrient-sensing organ that communicates systemic nutritional status to IPCs in the brain. Circulating amino acid uptake via the transporter slimfast (slif) and activation of the nutrient-responsive TORC1 pathway in the fat body promote the secretion of dIlps, particularly dIlp2, by IPCs^17^. Based on this, we hypothesized that Lst repression in EECs is not simply a local response to nutrient availability but is rather a gut response to low dIlp2 release by IPCs under nutritional stress. To test this, we impaired fat body nutrient sensing by overexpressing Tsc1*/2*, or by RNAi-mediated knockdown of *slif* or *Raptor* (Raptor being a TORC1 specific component^40^) using *Lpp-GAL4*^41^. Each manipulation, known to suppress fat body TORC1 signalling, further delayed development under malnutrition (Fig. S8A–C) and led to modest induction of *Lst* in the midgut despite the presence of microbial AAs (Fig. 8A–C). These findings suggest that TORC1 activity in the fat body is required to repress *Lst* expression in EECs in response to systemic cues. Because dIlp2 secretion by IPCs is a direct output of fat body nutrient sensing^17^, we next asked whether reduced dIlp2 production in IPCs is sufficient to trigger *Lst* expression.

**Figure 8.**
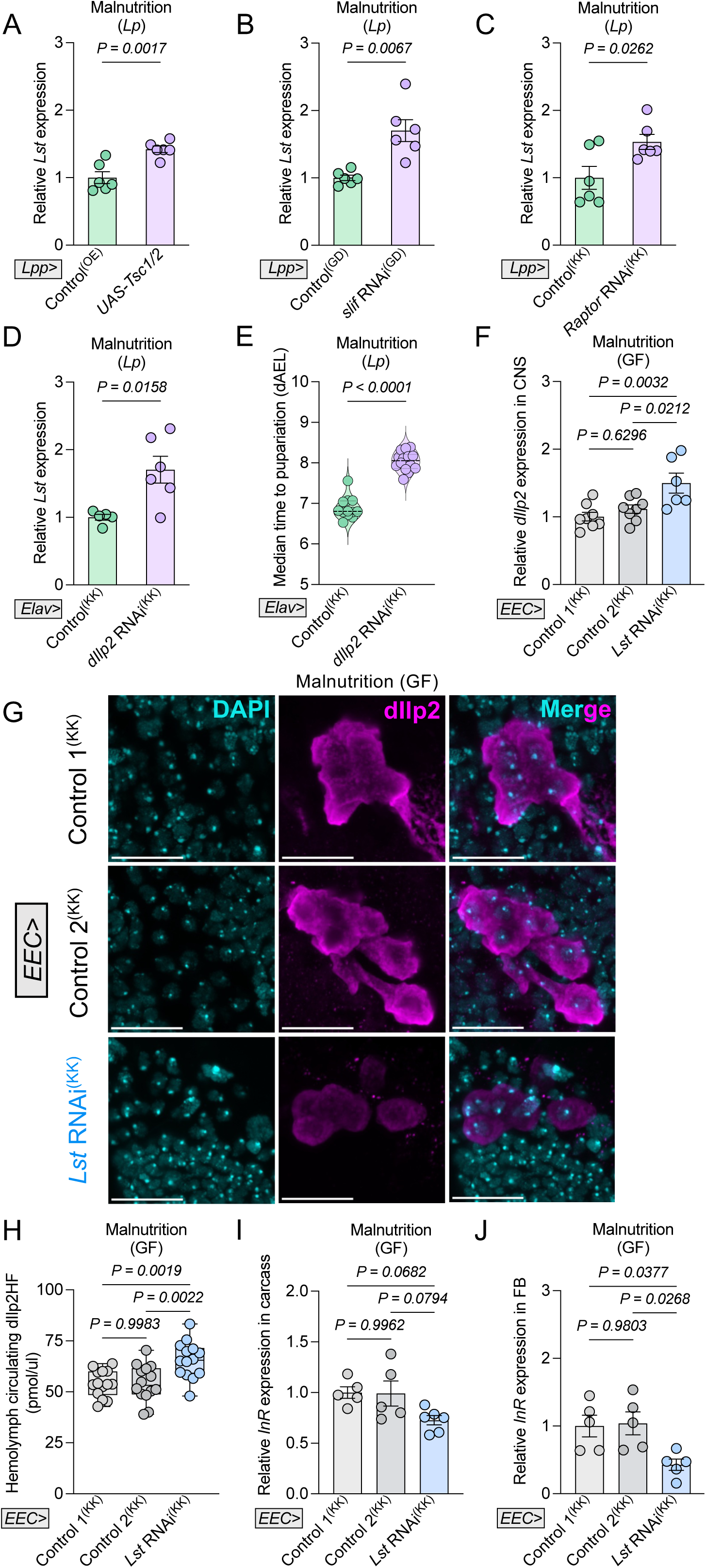
Lst is an enterokine involved in an inter-organ regulatory loop impacting circulating levels of dIlp2 in response to nutrient and microbiota status (A–D) Systemic TORC1 and insulin signalling in the fat body and central nervous system modulate gut *Lst* expression during malnutrition. Relative transcript levels of gut-derived *Lst* were analyzed by qRT-PCR in *Lp*-associated larvae with: (A) Overexpression of *Tsc1/2* in the fat body using the *Lpp-GAL4* (*Lpp>UAS-Tsc1/2*), compared to their respective overexpression controls; (B) Knockdown of *slif* in the fat body (*Lpp>UAS-slif RNAi*) compared to GD controls. (C) Knockdown of *Raptor* in the fat body (*Lpp>UAS-Raptor RNAi*) compared to KK controls; or (D) Knockdown of *dIlp2* in neurons including IPCs (*Elav>UAS-dIlp2* RNAi), compared to KK controls (*n* ≥ 5). (E) Median time to pupariation was measured for *Lp*-associated larvae with neuron-specific *dIlp2* knockdown (*Elav>UAS-dIlp2* RNAi) and their respective KK controls (*n* ≥ 12). (F–J) Under malnutrition, tissue responses were characterized in germ free larvae with EEC-specific *Lst* knockdown (*EEC>UAS-Lst RNAi*) and their respective KK controls. (F) Relative transcript levels of *dIlp2* in the brain analyzed by qRT-PCR. (G) Representative images of dIlp2 immunostaining in the brain; scale bar: 50 μm. (H) Circulating dIlp2(HF) levels in the hemolymph of heterozygous *Ilp2¹* gd2HF larvae measured by ELISA (Genotype: *EEC-GAL4/+; UAS-Lst RNAi/+; Ilp2¹*, gd2HF/+). (I, J) Relative transcript levels of *InR* analyzed by qRT-PCR in the carcass (I) and fat body (J). *n* ≥ 6 (F), *n* = 14 (H), *n* = 5 (I, J). In (A–D, F, I, J), data are presented as mean ± SEM. Statistical significance was determined using two-tailed unpaired *t* test (A, C, D), a two-tailed Welch’s *t* test (B, E) and one-way ANOVA with Tukey’s multiple comparisons test (F, H–G). Exact *P* values are indicated on the respective panels.

Pan-neuronal knockdown of *dIlp2* using *Elav-GAL4*^42^ significantly elevated *Lst* expression in the midgut (Fig. 8D) and delayed larval maturation into pupae (Fig. 8E), phenocopying the effects of fat body TORC1 inhibition.

Together, these results define a systemic nutrient-sensing loop linking dietary and microbial amino acid availability to Lst expression. In this loop, circulating amino acid are sensed by TORC1 in the fat body to promote dIlp2 release from IPCs, which maintains high IIS activity and suppresses Lst production in EECs. When circulating amino acid levels drop, reduced fat body TORC1 activity decreases dIlp2 release by the IPCs, leading to FoxO activation in Lst⁺ EECs and induction of Lst. This inter-organ circuit thus connects systemic amino acid sensing to gut endocrine output, coordinating developmental pace with environmental nutrient supply.

### Lst is an enterokine involved in an inter-organ feed-forward regulatory loop impacting circulating levels of dIlp2 in response to amino acids of dietary and microbial origins

Our data so far position Lst downstream of an inter-organ nutrient-sensing axis that links dietary and microbial amino acid availability to insulin signalling activity in the gut. However, the molecular consequence of Lst production and release by EECs under amino acid restriction remains unresolved. Given its endocrine nature and its previously described role in modulating dIlps release by IPCs in adult flies^27^, we hypothesized that Lst might act as an enterokine feeding back into the systemic nutrient-sensing circuit. Specifically, we asked whether Lst expression from EECs could influence the activity of IPCs in the brain, thereby modulating the production and release of dIlp2 in response to amino acids.

To test this, we examined dIlp2 expression at the transcript and protein levels in the central nervous system following targeted *Lst* knockdown in EECs. This led to a clear increase in *dIlp2* transcript levels in the brain (Fig. 8F) and a marked decrease in dIlp2 staining in IPCs (Fig. 8G), suggesting that Lst from EECs negatively regulates *dIlp2* expression in the brain and stimulates dIlp2 retention in IPCs. We next assessed circulating dIlp2 levels under similar conditions. Strikingly, repression of *Lst*, either in Lst^+^ cells (Fig. S8D) or in EECs (Fig. 8H) of malnourished germ-free animals, resulted in a significant increase in hemolymph dIlp2 concentration with levels approaching those observed in malnourished animals mono-associated with *Lp* (Fig. S8E). Accordingly, the transcript levels of *InR*, a well-established (repressed) transcriptional readout of IIS showed a trend to a reduction in the carcass (Fig. 8I) and significantly downregulated in the fat body (Fig. 8J), upon *Lst* knockdown in EECs, further confirming that the knockdown of gut-derived *Lst* leads to elevated systemic insulin activity across peripheral tissues.

Taken together, these findings establish Lst as an amino acid-responsive enterokine that contributes to an inter-organ regulatory loop engaged under malnutrition. By modulating dIlp2 production and release from brain IPCs, Lst helps adjust developmental progression at the systemic level in response to fluctuations in dietary and microbial amino acid supplies.

## Discussion

By studying *Drosophila* larval development under conditions of severe amino acid restriction, achieved through either extremely low yeast diet or low-yeast diet coupled with microbiota depletion, we uncover a critical inter-organ communication axis involving nutrient assimilation by enterocytes, nutrient sensing in fat body, dIlp production in IPCs, and signalling feedbacks on EECs via IIS. This axis leads to the regulation of Lst, an amino acid-responsive enterokine secreted by a small population of anterior midgut EECs. Lst acts as a potent inhibitor of dIlp2 circulating levels, a key insulin-like peptide driving systemic developmental progression in larvae. Our findings reveal a novel inhibitory loop that couples availability of amino acid from dietary or microbial origins to developmental pace through the downregulation of dIlp2 expression and secretion.

In their natural environment such as decaying fruit, *Drosophila* larvae frequently face protein content fluctuations. The microbiome with its ability to produce amino acid serves as a critical nutritional buffer in these contexts, and the Lst-insulin feedback loop allows larvae to tightly modulate their developmental trajectory in response to nutritional stress, thereby preserving viability and ensuring successful maturation to the adult stage (Fig. 9). In this way, Lst functions as a critical adaptive signal that links nutrient sensing to whole-body growth and maturation control, a key determinant of survival under nutritionally challenging conditions.

**Figure 9.**
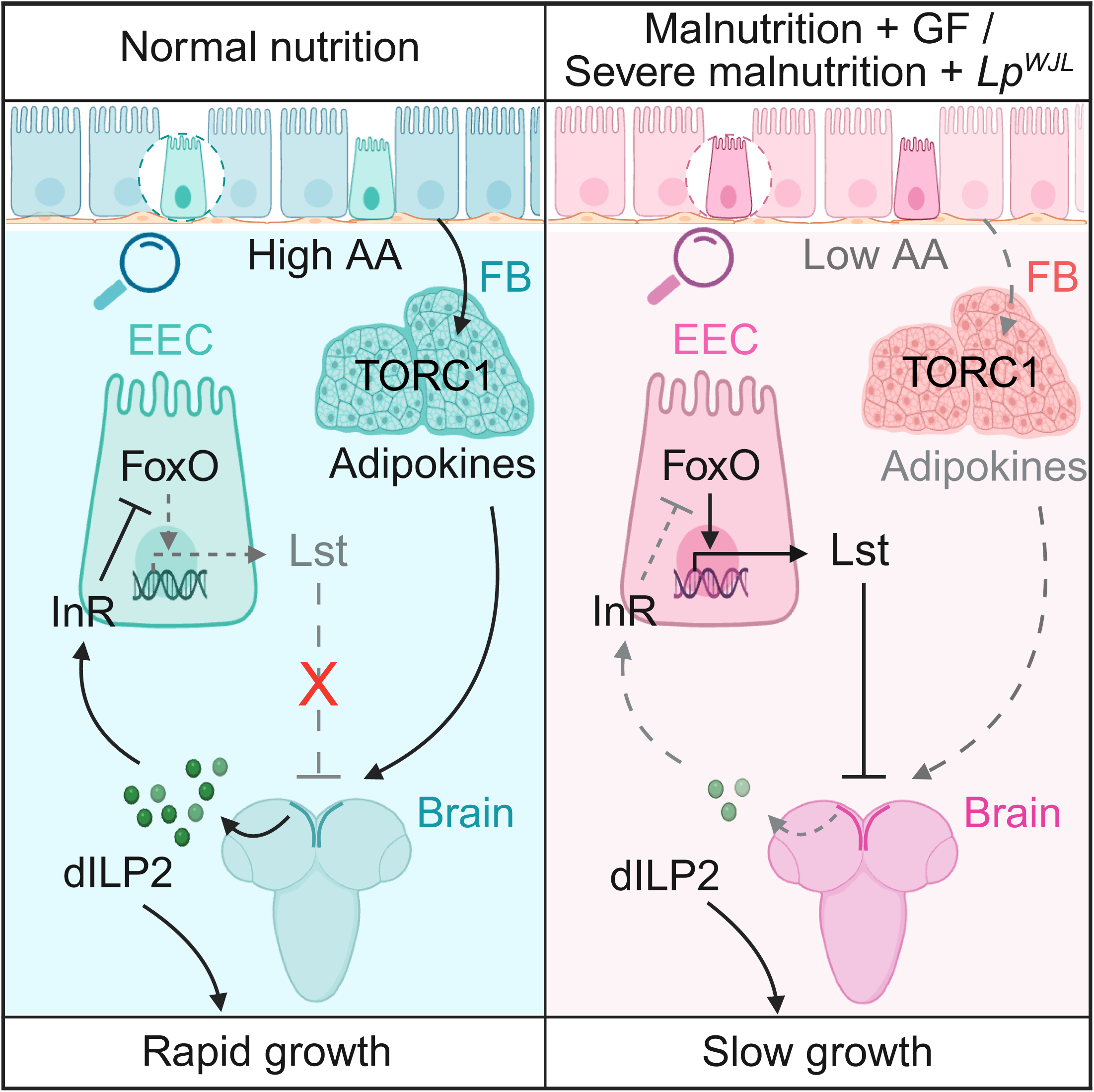
Schematic model illustrating the gut–brain axis regulating larval adaptation to nutritional stress Compared with normal fed conditions, undernourished germ-free (GF) status or severe malnutrition associated with *Lp^WJL^* induces *Lst* expression in a small population of midgut EECs. Once induced, gut-derived Lst acts systemically to suppress the circulating levels of dIlp2. This EEC-specific Lst up-regulation is triggered by reduced amino acid availability through a multi-organ relay involving the fat body and IPCs in the brain. Consequently, Lst orchestrates a feedback control loop that temporarily attenuates developmental progression upon sensing systemic nutrient shortages. This bidirectional gut–brain axis ultimately enables *Drosophila* larvae to preserve viability under severe nutritional stress.

These findings build upon the growing understanding that the gut is more than a digestive organ, it also functions as a central endocrine hub capable of orchestrating systemic physiological responses^43^. Our results identify Lst as a key molecular mediator of this gut-endocrine function, acting in a nutrient-sensitive and gut-derived manner to systemically modulate insulin signalling and developmental pace. Notably, while contributions from other cell populations cannot be entirely excluded, we find that only less than 10 Lst^+^ EECs located in the anterior midgut are sufficient to exert this regulatory function. This small and spatially restricted cell population has a disproportionately large impact on systemic physiology, acting as a critical node in the relay of nutritional information to the IPCs another small group of cells with a large impact on systemic regulation of developmental pace.

While Lst was originally described in corpora cardiaca cells as a sugar-responsive regulator of insulin secretion^27^, our results highlight a distinct, juvenile-specific mechanism of Lst regulation and function. In larvae, gut-derived Lst responds to availability of amino acid from dietary and/or microbial origins, whereas corpora cardiaca-derived Lst does not exhibit regulation in this nutritional context. This suggests that the larval midgut EECs are the primary source of nutritionally responsive Lst in early life. In larvae gut-derived Lst only mildly responded to full dietary sugar depletion (data not shown) while in adults, Lst+ EECs were not identified^27^ (data not shown) and corpora cardiaca-derived Lst responds to sugar rather than protein, similarly to adult IPCs. In contrast, larval IPCs are more responsive to amino acids, major determinants of somatic growth and maturation, suggesting a developmental switch in both nutrient sensing and hormonal regulation of IPCs function aligned with a juvenile physiological demand high in proteins while adult maintenance mainly rely on energy. These data further emphasize the unique contribution of gut-derived Lst in the larval stage.

How Lst impinges on IPCs function to modulate dIlp2 production remains an important question. Notably, Lst^+^ EECs are positioned on the basal side of the larval midgut epithelium, consistent with a model in which Lst is secreted basally into the hemolymph to act systemically. Our current data confirm its long-range endocrine action regulating dIlp2 circulating levels, but the identity of the receptor mediating Lst function in IPCs is still unresolved. While the Pyrokinin1 receptor (Pk1-R) was previously proposed as a candidate^27^, our tests in larvae did not support its involvement (Fig. S8F–H). Identifying the receptor for Lst will be key to fully elucidating its mechanism of action.

Why would such a complex inter-organ relay, gut to fat body (via circulating amino acids), fat body to IPCs (via adipokines) and back to gut (via dIlps), evolve to regulate Lst secretion? One possible explanation is that Lst being a potent inhibitor of dIlp2 release, it needs to be tightly regulated. Such a multi-step regulatory architecture may ensure that Lst production is fine-tuned to regulate organismal growth in response to environmental cues and only deployed under sufficiently severe nutrient stress, thereby preserving dIlp2-driven growth unless absolutely necessary. This regulatory loop may serve as a threshold switch, engaged only when amino acids provision falls below critical levels to ensure survival. In this context, the basal side of the gut epithelium, positioned to receive systemic hormonal and metabolic cues, could act as both a sensor and amplifier of whole-body nutrient status, modulating enterokine secretion accordingly. The small group of Lst^+^ EECs may indeed function as a signalling center, analogous to the IPCs, which also comprise only a few pairs of neurosecretory cells yet exert profound systemic effects on growth and metabolism. At the same time, such a centralized signalling role does not exclude the existence of additional local sensing activities of Lst^+^ EECs that may not be captured in the present study. However, the basal location of Lst^+^ EECs is more aligned with a sensing and signalling activity from and to the “Internal milieu” rather than a sensing and signalling activity of the intestinal luminal content.

Taken together, our findings establish the larval gut as a nutrient-sensitive endocrine organ capable of coordinating developmental plasticity. The identification of Lst as an enterokine inhibiting dIlp2 secretion reveals a new regulatory logic by which nutrient availability is translated into hormonal control of developmental pace and ultimately survival. Our study expands the conceptual framework of decretins, hormones that suppress insulin production in the context of adult sugar metabolism, into the realm of developmental control. In *Drosophila*, Lst exemplifies how a nutrient-sensitive enterokine, a gut-derived hormone, can shape developmental trajectories. Notably, while the decretin concept has been difficult to confirm in mammals in the context of insulin regulation by pancreatic beta cells^44^, our findings raise the possibility that analogous enterokines may act instead on tissue responsiveness to somatotropic hormones such as growth hormone or directly on hepatic IGF1 production. Thus, Lst may point to a broader, evolutionarily conserved or convergent mechanisms in which the gut serves as an endocrine modulator of systemic adaptation at the juvenile stage by regulating IGF1-mediated developmental responses in a nutrient-dependent manner.

## Methods

### Fly stocks

*y,w* (gift from Bruno Lemaitre), *AKH-GAL4* (BDSC#25684), *EEC-GAL4* (BDSC#84614), *voilà-GAL4* (BDSC#84276), *mex-GAL4* (gift from Irene Miguel-Aliaga), *Lst-GAL4* (gift from Seung K Kim), *Lpp-GAL4* (gift from Alexandre Djiane), *Elav-GAL4* (BDSC#458), *Lst^1^* (gift from Seung K Kim), *Lst^CRIMIC^* (BDSC#91330), *dIlp2^1^-gd2HF* (gift from Pierre Léopold), *UAS-GFP* (cytoplasmic GFP; gift from Jonathan Enriquez), *Lst* RNAi^KK^ (VDRC#106861), *Lst* RNAi^TRiP^ (BDSC#60400), *Gcn2* RNAi^TRiP^ (BDSC#67215), *FoxO* RNAi^KK^ (VDRC#107786), *slif* RNAi^GD^ (VDRC#45587), *Tsc1* RNAi^KK^ (VDRC#110811), *InR* RNAi^GD^ (VDRC#992), *Raptor* RNAi^KK^ (VDRC#106491), *Pten* RNAi^KK^ (VDRC#101475), *dIlp2* RNAi^KK^ (VDRC#102158), *PK1-R* RNAi^KK^ (VDRC#101115), *PK1-R* RNAi^TRiP^ (BDSC#27539), *PK1-R* RNAi^SH^ (VDRC#330176), GD control (VDRC#60000), KK control 1 (VDRC#60102), KK control 2 (designated as KK control) (VDRC#60103), TRiP control 1 (BDSC #36304), TRiP control 2 (designated as TRiP control) (BDSC#35785), SH control (VDRC#60201), *UAS-Lst* (gift from Seung K Kim), *UAS-FoxO* (BL#9575), *UAS-InR^DN^* (BL#8253), OE control (BL#35786). To eliminate potential genetic background confounding, all experimental lines (RNAi or overexpression) and their respective controls were crossed to the identical GAL4 driver line, ensuring a highly consistent genetic architecture across all conditions.

### Fly husbandry

Conventional fly stocks were raised on a yeast/cornmeal medium (7.14 g/L of agar, 50 g/L of inactivated yeast, 80 g/L of cornmeal, 4 mL/L of propionic acid, and 5.2 g/L of Methyl 4-hydroxybenzoate sodium salt) at 25°C under a 12:12 hour light-dark cycle. germ free flies were generated as previously described^45^ and raised on a yeast/cornmeal diet supplemented with four antibiotics (50 µg/mL of ampicillin, 5 µg/mL of erythromycin, 50 µg/mL of kanamycin, and 10 µg/mL of tetracycline). For the experimental malnutrition condition, the concentration of inactivated yeast in the medium was reduced to either 7 or 3 g/L (malnutrition or severe malnutrition, respectively).

### Holidic diets

Holidic Diets (HDs) were prepared based on previously described FLYAA recipes^33^, with minor modifications to the amino acid composition. Specifically, the HAAHD contained a total amino acid concentration of 21.38g/L, while the LAAHD contained 8.55g/L (representing a 60% reduction in the concentration of all AAs compared to the HAAHD). More specific HD compositions were generated by further manipulating the amino acid content of the initial LAAHD and HAAHD formulations. Specifically, the six essential amino acids produced by *Lp* (histidine, lysine, methionine, phenylalanine, threonine, and asparagine) were either reduced by 60% from the HAAHD level or increased by 2.5 times the LAAHD concentration. To prevent the inhibition of bacterial growth in gnotobiotic experiments, chemical preservatives such as propionic acid or nipagin were intentionally omitted from all HD formulations. To ensure sterile conditions for experiments, all tubes were UV-treated, egg-laying cages were autoclaved, and all diet solutions were either filter-sterilized or autoclaved as appropriate. Freshly prepared HDs were stored at 4°C and used within one week of preparation.

### Bacterial culture

*Lp^WJL^* was cultured in MRS Broth (Carl Roth) overnight without shaking at 37°C. *Ap^WJL^* was cultured in 10 mL of Mannitol Broth—comprising 3 g/L Bacto peptone, 5 g/L yeast extract (both from Becton Dickinson), and 25 g/L D-mannitol (Carl Roth)—in a 50-mL flask at 30°C with shaking at 180 rpm for 24 h. After centrifugation, the supernatant was discarded and the bacterial pellet was resuspended in sterile PBS for fly food inoculation at a final concentration of 10^8^ CFU/tube for *Lp^WJL^* and 10^5^ CFU/tube for *Ap^WJL^*.

### Developmental timing experiments

To establish germ free and mono-associated conditions, germ free adult flies were allowed to lay eggs overnight in sterile breeding cages containing fly food medium. Axenic embryos were collected and transferred to individual fly food tubes. Each tube was then inoculated with 200 µL of either sterile PBS for the germ-free condition or 200 µL of bacteria resuspended in sterile PBS, for mono-association experiments. Unless otherwise specified, all fly lines were reared at a standard temperature of 25°C under a 12:12 hour light-dark cycle. Larval development and growth rate were assessed by monitoring pupariation time. The number of pupae in each tube was counted daily until the emergence of the last pupa.

The median time required for larvae to reach pupariation after egg laying was used as a quantitative measure of the larval growth rate. Each data point represents the median time to pupariation for one tube of pupae (40 ± 15 pupae).

### Larval size measurements

For larval size measurements, *Drosophila* larvae were collected and washed with PBS. The larvae were then transferred to a beaker containing PBS and euthanized using a brief heat treatment (60°C). Subsequently, the larvae were positioned for imaging and photographed using a Leica M205FA Stereomicroscope. Larval longitudinal size was quantitatively measured from the resulting images using the Fiji software (Version 1.54r).

### Food intake experiments

Stage-(L3) and Size-(3.5mm) matched larvae were collected and placed on the low-yeast diet plates supplemented with 0.8% Erioglaucine disodium salt (Sigma-Aldrich, #861146) for either 1 hour or 2 hours. Following the feeding period, larvae were washed with distilled PBS to remove residual surface dye. Seven larvae were then pooled in a 2 mL Eppendorf tube containing beads and 500 µL PBS. Larvae were homogenized using a Precellys 24 tissue homogenizer (Bertin Technologies) with two cycles of 30 s at 6000 rpm, separated by a 30 s pause. Homogenates were then centrifuged for 60 s at 10,000 × *g*. The absorbance of the resulting supernatant was measured at 629 nm using a microplate reader (BMG LABTECH).

### Quantitative real-time PCR

Total RNA was extracted from dissected tissues of stage-and size-matched larvae using the NucleoSpin RNA (MACHEREY-NAGEL, #740955) or NucleoSpin RNA XS (MACHEREY-NAGEL, #740902). RNA was reverse-transcribed into cDNA using the SensiFAST cDNA Synthesis Kit (Bioline, #BIO-65054). Quantitative real-time PCR (qPCR) was performed using the No ROX SYBR 2X MasterMix Blue dTTP (Takyon, #UF-NSMT-B0701) on a CFX96 Real-Time System (BIO-RAD). All expression levels were normalized to two reference genes, *RpL32* and *Tubulin* using delta-delta-Ct method. The following primers (5’–3’) were used for qPCR assays: *Tubulin* qF TGTCGCGTGTGAAACACTTC, *Tubulin* qR AGCAGGCGTTTCCAATCTG; *Rpl-32* qF ATGCTAAGCTGTCGCACAAATG, *Rpl-32* qR GTTCGATCCGTAACCGATGT; *Lst* qF ATAGTCTGCGATCCAAGCCG, *Lst* qR CTGCAGACGGTTCAGATCGT; *InR* qF AAGCGTGGGAAAATTAAGATGGA, *InR* qR GGCTGTCAACTGCTTCTACTG; *dIlp2* qF CGAGGTGCTGAGTATGGTGTG, *dIlp2* qR CCCCCAAGATAGCTCCCAGGA.

### Hemolymph Collection

A 20 μL filtered pipette tip and a 10 μL filtered pipette tip were each cut at an angle. The modified 20 μL tip was placed inside a 2 mL Eppendorf tube, and the modified 10 μL tip was then inserted into the wider opening of the 20 μL tip, creating a nested collection system.

Stage-and Size-matched *Drosophila* larvae were washed with PBS, and excess surface moisture was carefully removed using a paper towel. Individual larvae were transferred to the inner wall of the 10 μL pipette tip, where they were gently torn open from the anterior end to release hemolymph. Hemolymph was collected into the 2 mL Eppendorf tube by centrifuging the assembly at 10,000 × *g* for 1 min at 4°C. The collected hemolymph was immediately flash-frozen in liquid nitrogen and subsequently stored at –80°C. To minimize protein degradation, all steps of the hemolymph collection procedure were performed on ice.

### dIlp2-HF ELISA

ELISA was performed using Nunc-Immuno modules (ThermoFisher Scientific, #468667). Each well was initially coated with 100 μL of anti-FLAG antibody (Sigma-Aldrich, #F1804), diluted to a concentration of 5 μg/mL in 0.2 M sodium carbonate/bicarbonate buffer (pH 9.4). The modules were incubated overnight at 4°C to allow for antibody binding. Following the coating step, the plate was washed three times with 0.2% PBST (PBS containing 0.2% Tween 20). Non-specific binding sites were then blocked by adding 350 μL of blocking buffer (PBS containing 2% BSA (ThermoFisher Scientific, #37525) to each well and incubating for 2 hours at room temperature. After blocking, the plate was washed three times with 0.2% PBST. Subsequently, 1 μL of either collected hemolymph sample or synthetic FLAG(GS)HA peptide standard was mixed with 50 μL of detection antibody solution. The detection antibody solution consisted of anti-HA-Peroxidase 3F10 antibody (Roche, #12013819001) at a concentration of 25 ng/mL in 0.1% PBSTx (PBS containing 0.1% Triton X-100). This mixture was then added to the appropriate wells. The wells were sealed, and the plate was incubated overnight at 4°C. The solution was removed from the wells by aspiration, and the plate was washed thoroughly six times with 0.2% PBST. For color development, 100 μL of 1-Step Ultra TMB ELISA Substrate (ThermoFisher Scientific, #34028) was added to each well. The plate was incubated on a rotary shaker for 30 min at room temperature to facilitate the enzymatic color reaction. The reaction was terminated by adding 100 μL of 2 M sulfuric acid to each well. The absorbance at 450 nm was immediately measured using a microplate reader (BMG LABTECH).

### Anti-Lst antibody generation

A rabbit polyclonal anti-Lst serum was produced by Pacific Immunology Inc. A 15-amino acid peptide (Cys-AIVFRPLFVYKQQEI) was synthesized and used for rabbit immunization. Lst antibody was purified by affinity chromatography from a 25 mL serum pool collected from two rabbits.

### Immunofluorescence

Fresh tissues from stage-and size-matched larvae were dissected in cold PBS and fixed in 4% paraformaldehyde (PFA) (16% PFA (ThermoFisher Scientific, #28906) diluted in PBS) for 60 min with gentle agitation. After fixation, the tissues were permeabilized by washing 3 times with 0.1% PBSTx and incubated with 3% goat serum (Sigma-Aldrich, #G9023) in 0.1% PBSTx at RT with shaking. The tissues were then incubated with primary antibodies overnight at 4°C. After the primary antibody solution was removed, the tissues were washed 5 times with 0.1% PBSTx and further incubated with secondary antibodies diluted in 0.1% PBSTx for 2h at RT. After 5 washes with 0.1% PBSTx, the samples were mounted in ROTI Mount FluorCare DAPI (Carl Roth, #HP20.1). To reduce the strong background of Lst staining, the Lst antibody was pre-cleared using posterior larval midguts and re-collected for the experiments. Details of the antibodies used in this study are listed below: chicken anti-GFP (Abcam, #ab13970, 1:1000), mouse anti-Prospero (MR1A) (DSHB, 1:10), rabbit anti-Lst (Custom-made, 1:50), rabbit anti-dIlp2 (gift from Jan A Veenstra, 1:500), rabbit anti-Phospho-Drosophila Akt (Ser505) (Cell Signalling Technology, #4054, 1:200), Alexa Fluor 488-conjugated goat anti-chicken (ThermoFisher Scientific, #A11039, 1:200), Alexa Fluor 555-conjugated goat anti-rabbit (ThermoFisher Scientific, #A21428, 1:200), and Alexa Fluor 647-conjugated goat anti-mouse (ThermoFisher Scientific, #A21235, 1:200).

Samples were imaged using a Leica SP8 confocal microscope. To compare Lst, dIlp2, and pAKT protein levels, all samples were imaged using identical acquisition settings. Image processing was performed using Fiji software, and 3D reconstructions were generated using the ImageJ 3D Viewer plugin (Version 1.54r).

## Statistical analyses

All statistical analyses were conducted using GraphPad Prism 10 software (Version 10.6.1). Detailed statistical methods are indicated in the figure legends. Normality and lognormality tests were conducted. Statistical significance was analyzed using *t* tests, one-way ANOVA, and two-way ANOVA (employing the Geisser-Greenhouse correction). Detailed statistical methods are indicated in the figure legends.

## Acknowledgements

The authors would like to thank Rénald Delanoue, Pierre Léopold, Gilles Storelli, Pavel Melentev, Manon Picquenot, Juliette Mendes, Houssam Akherraz, Ziyan Nie, Julie Carnesecchi and Pedro Borges Pinto for help and discussions; Seung Kim, Pierre Leopold, Irene Miguel-Aliaga, Alexandre Djiane, Bruno Lemaitre and Jonathan Enriquez for sharing reagents; the IGFL imaging facility; the PLATIM and Arthro-Tools platforms of the SFR Biosciences Lyon Gerland (UAR3444/US8) for providing Imaging and *Drosophila* facilities as well as the Bloomington Drosophila Stock Center (BDSC) and Vienna Drosophila Ressource Center (VDRC) for fly stocks. This work was funded by ANR-23-CE14-0034 (to F.L), FRM DEQ20180839196 (to F.L), UCBL BQR (to C.I.R) and FINOVI AO12-19-Projet SNI (to C.I.R). LB was funded by a Chinese Government Scholarship.

## Declaration of generative AI and AI-assisted technologies in the writing process

During the preparation of this work, the authors used ChatGPT in order to polish the language of the manuscript. After using this tool, the authors reviewed and edited the content as needed and take full responsibility for the content of the publication.

## Movie S1 (related to Figure 2)

The spatial distribution of *Lst*-expressing EECs in the gut was visualized by 3D reconstruction using the *Lst-GAL4>GFP* reporter line. DAPI (nuclei) is shown in gray and GFP (*Lst*-expressing EECs) in green.

---

**Figure S1.**
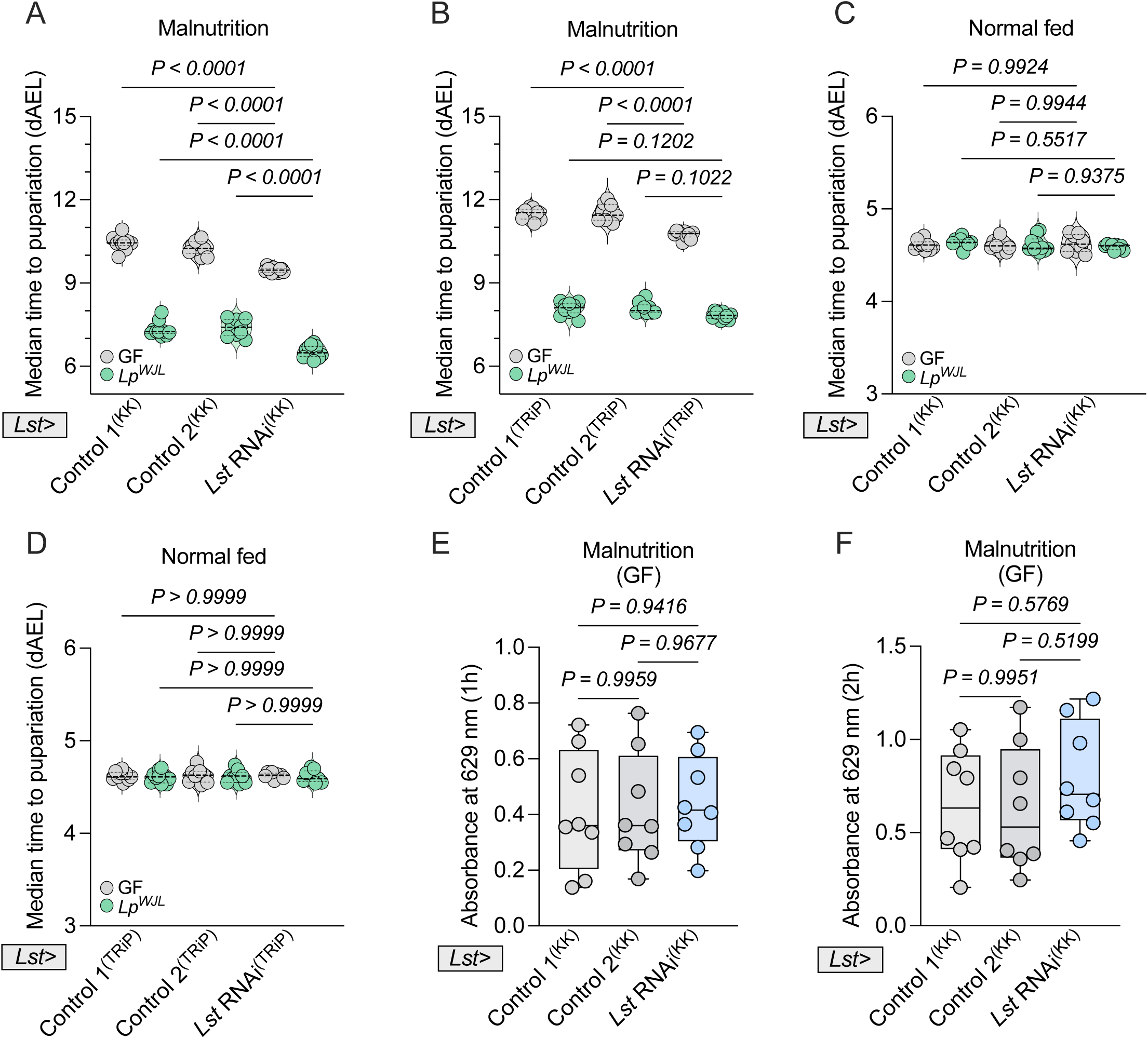
(related to Figure 1). Knockdown of *Lst* accelerates larval systemic growth specifically under malnutrition (A–D) Median time to pupariation measured for *Lst-GAL4* mediated *Lst* RNAi larvae and their respective controls. Assessments were performed under malnutrition (A, B) or normal fed conditions (C, D), utilizing either the KK RNAi library (A, C) or the TRiP RNAi library (B, D) (*n* ≥ 9). (E, F) Food intake quantification for undernourished germ-free larvae with *Lst-GAL4* driven *Lst* knockdown (*Lst>UAS-Lst* RNAi) and their corresponding KK control larvae after 1 hour (E) and 2 hours (F) of feeding (*n* = 8). Statistical significance was determined using a two-way ANOVA with Tukey’s multiple comparisons test (A–D) or a one-way ANOVA with Tukey’s multiple comparisons test (E, F). Exact *P* values are indicated on the respective panels.

**Figure S2.**
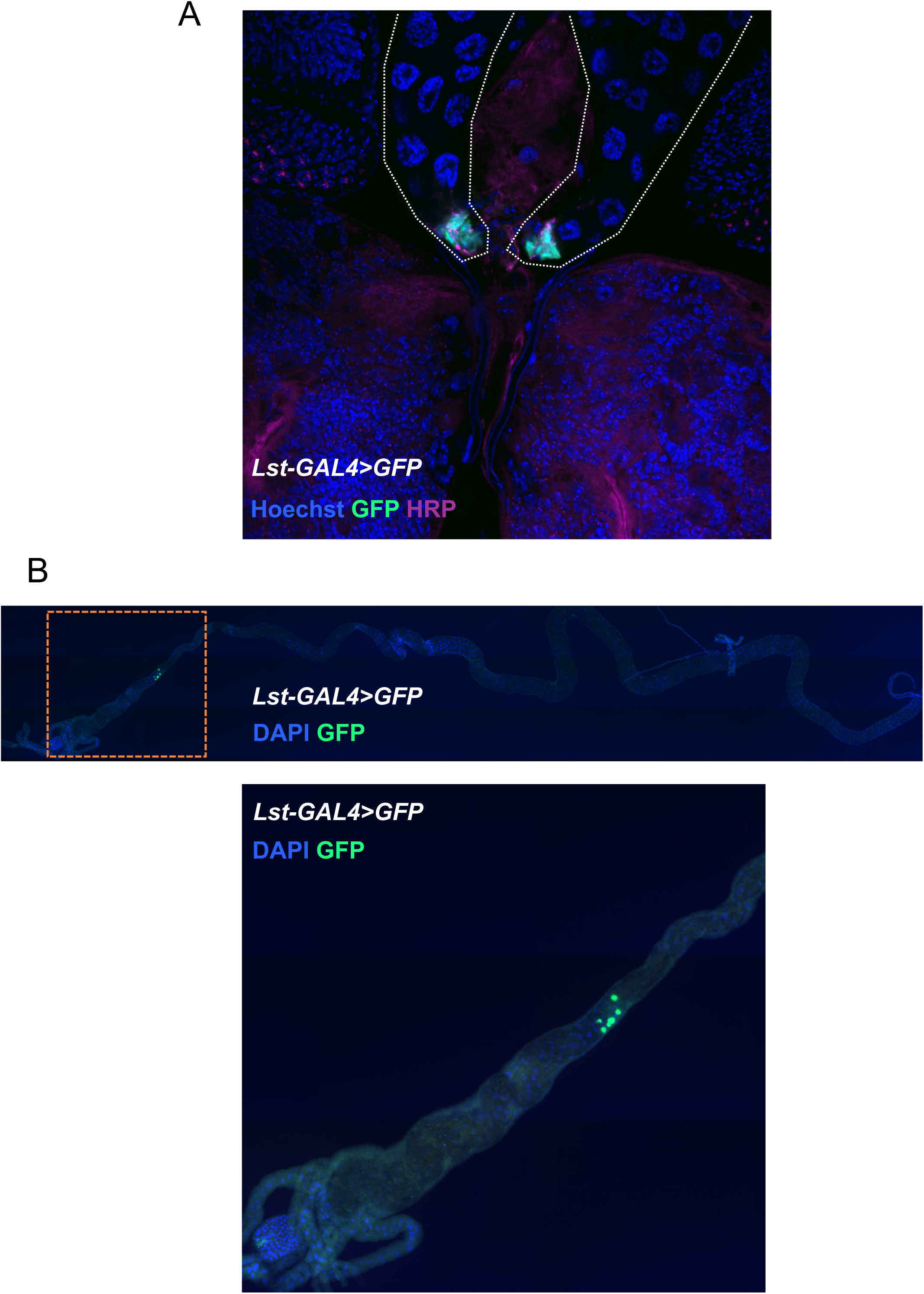
(related to Figure 2). *Lst* expression patterns in *Drosophila* larvae (A, B) Spatial distribution of *Lst* expression revealed by the *Lst-GAL4>GFP* reporter line under germ free and malnourished conditions. (A) The reporter line driving GFP signals in corpora cardiaca (CC); the ring gland is outlined by dashed lines, and the corpora cardiaca is localized at the base of the ring gland structure. Anti-HRP (horseradish peroxidase) staining was utilized to label the secretory neurons. (B) GFP-positive EECs localized in the midgut region anterior to the acidic zone.

**Figure S3.**
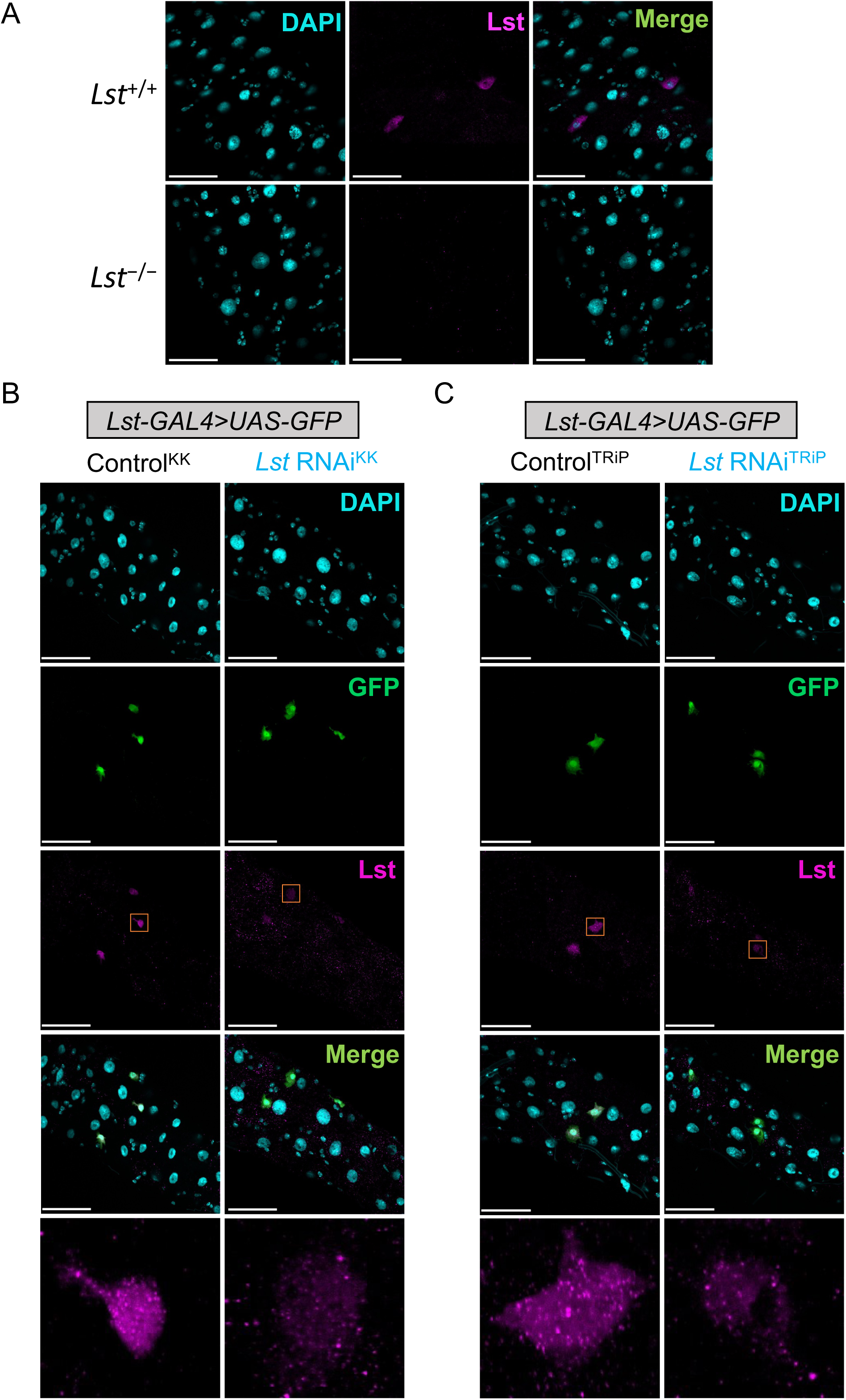
(related to. **Figure 2). Validation of Lst antibody specificity and genetic tools** (A–C) Midgut anti-Lst immunostaining in germ-free (GF) malnourished larvae. (A) The Lst peptide was visualized in the midguts of *yw* control larvae and *Lst*^−/−^ null mutant larvae. (B, C) Validation of *Lst* RNAi lines via Lst immunostaining of *Lst-GAL4*-mediated *Lst* RNAi larvae, utilizing either the KK RNAi library (B) or the TRiP RNAi library (C). Scale bar: 50 μm.

**Figure S4.**
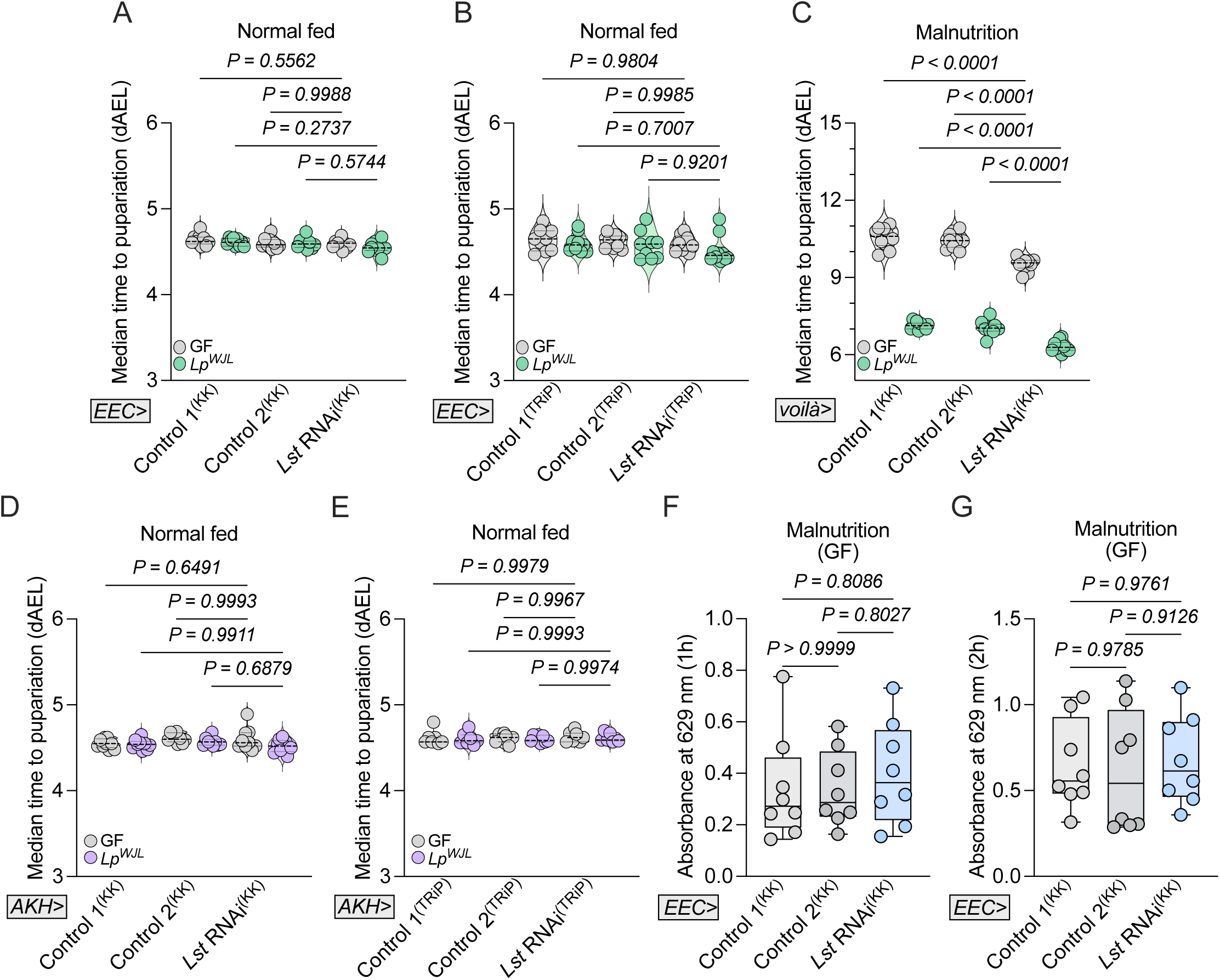
(related to Figure 4). Tissue-specific *Lst* knockdown reveals tissue-dependent roles in larval developmental timing (A, B) Median time to pupariation was measured under normal fed conditions in larvae with *Lst* knockdown specifically in EECs (*EEC>UAS-Lst* RNAi), compared to their respective controls (*n* ≥ 9). Data are shown for the KK RNAi line (A) and the TRiP RNAi line (B). (C) Median time to pupariation was measured under malnutrition in larvae with EEC-specific *Lst* knockdown driven by the *voilà-GAL4* (*voilà>UAS-Lst* RNAi), compared to KK control larvae. (*n* = 10). (D, E) Median time to pupariation was measured under normal fed conditions in larvae with *Lst* knockdown in the corpora cardiaca (*AKH>UAS-Lst* RNAi), compared to their respective controls (*n* ≥ 7). Data are shown for the KK RNAi line (D) and the TRiP RNAi line (E). (F, G) Food intake was quantified over 1 hour (F) and 2 hours (G) in undernourished germ-free larvae with EEC-specific *Lst* knockdown (*EEC > UAS-Lst* RNAi), compared to KK control larvae. (*n* = 8). Statistical significance was determined using a two-way ANOVA with Tukey’s multiple comparisons test (A–E) or a one-way ANOVA with Tukey’s multiple comparisons test (F, G). Exact *P* values are indicated on the respective panels.

**Figure S5.**
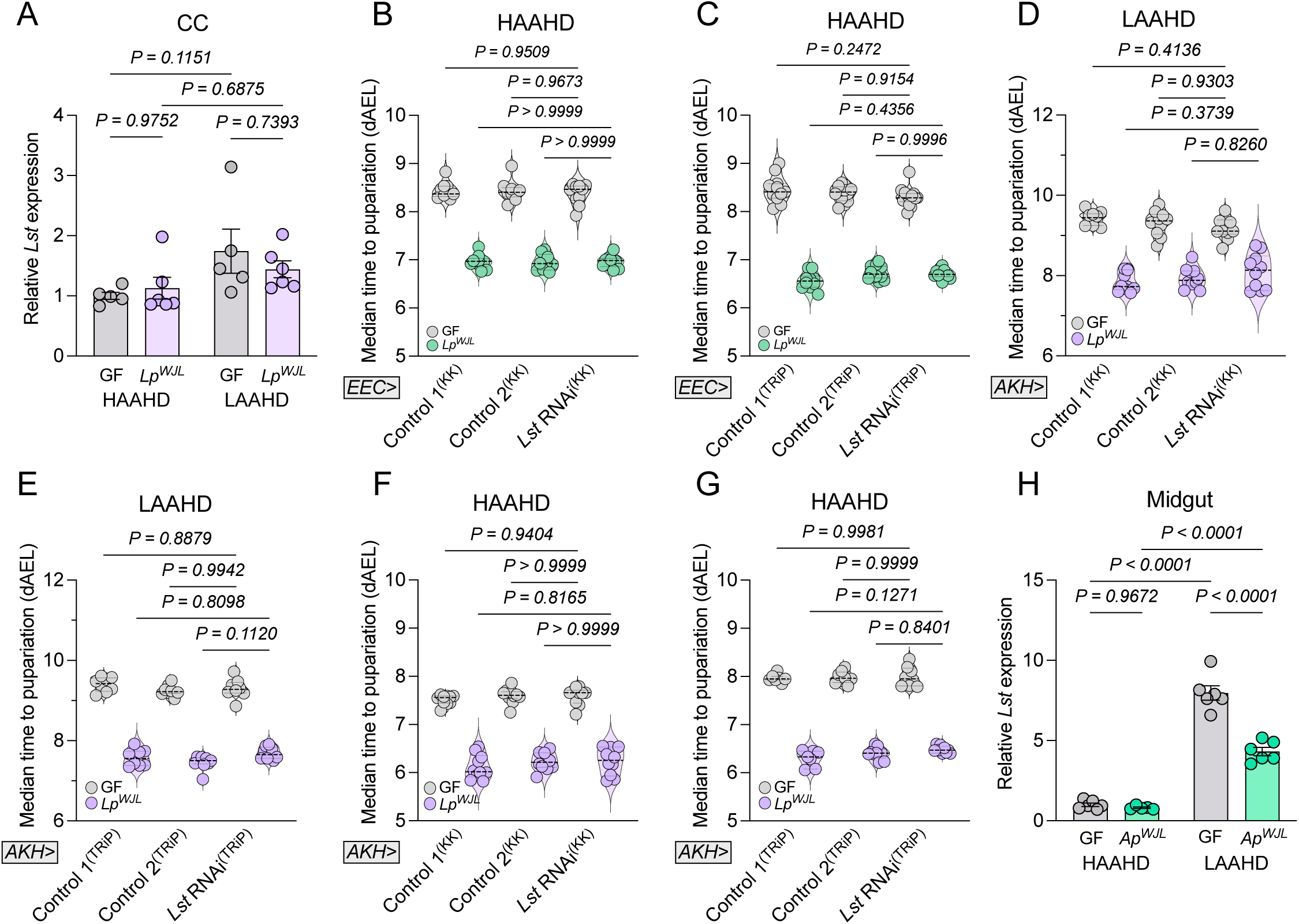
(related to Figure 5). The expression of corpora cardiaca-derived Lst is insensitive to dietary AAs at the larval stage (A, H) Relative transcript levels of corpora cardica-derived *Lst* (A) or gut-derived *Lst* (H) analyzed by qRT-PCR in GF, *Lp*-associated (A), or *Ap*-associated (H) larvae reared under HAAHD or LAAHD conditions (*n* ≥ 5). (B, C) Median time to pupariation was measured under in larvae reared under HAAHD conditions with EEC-specific *Lst* knockdown (*EEC>UAS-Lst* RNAi), compared to their respective controls (*n* ≥ 9). Data are shown for the KK RNAi line (B) and the TRIP RNAi line (C) (D–G) Median time to pupariation was measured in larvae with corpora cardiaca-specific *Lst* knockdown (*AKH>UAS-Lst* RNAi), compared to their respective controls (*n* ≥ 7), reared either under LAAHD conditions (D, E) or under HAAHD conditions (F, G). Data are shown for the KK RNAi line (D, F) and the TRiP RNAi line (E, G). In (A, H), data are presented as mean ± SEM. Statistical significance was determined using a two-way ANOVA with Tukey’s multiple comparisons test (A–H). Exact *P* values are indicated on the respective panels.

**Figure S6.**
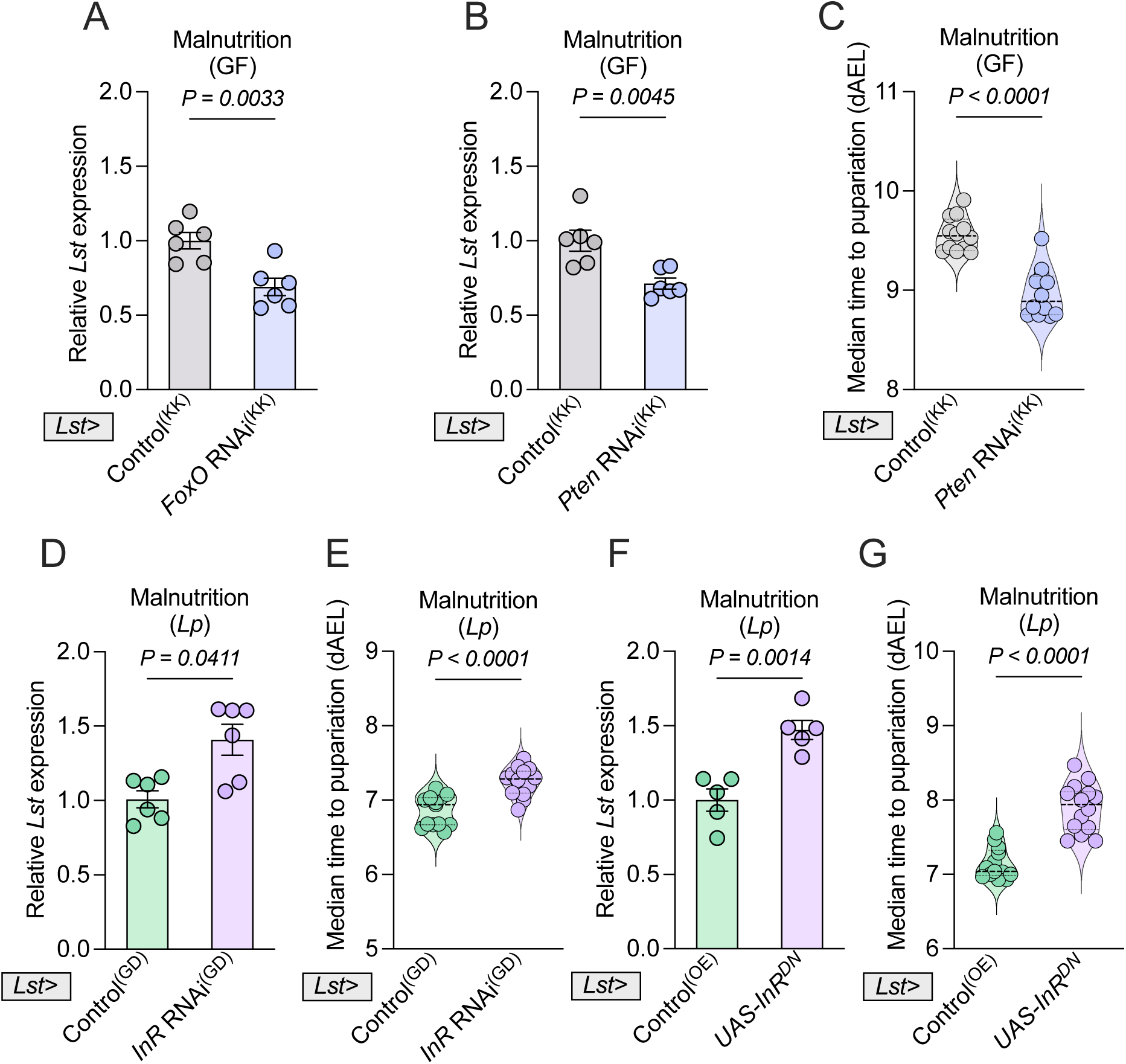
(related to Figure 7). *Lst* induction in EECs is controlled by reduced systemic insulin signalling via FoxO. (A) Under malnutrition, relative transcript levels of gut-derived *Lst* analyzed by qRT-PCR in germ free larvae with *FoxO* knockdown in *Lst*-expressing cells (*Lst>UAS-FoxO* RNAi), compared to their respective KK controls (*n* = 6). (B, C) Relative transcript levels of gut-derived *Lst* analyzed by qRT-PCR (B) and median time to pupariation measured (C) under malnutrition for germ free larvae with *Pten* knockdown in *Lst*-expressing cells (*Lst>UAS-Pten* RNAi), compared to their respective KK controls. *n* = 6 (B), *n* = 12 (C). (D–G) Under malnutrition, relative transcript levels of gut-derived *Lst* analyzed by qRT-PCR (D, F) and median time to pupariation measured (E, G) for *Lp*-associated larvae with *InR* knockdown in *Lst*-expressing cells (*Lst>UAS-InR* RNAi) and their respective GD controls (D, E), or with specific overexpression of dominant-negative *InR* in *Lst*-expressing cells (*Lst>UAS-InR^DN^*) and their corresponding overexpression controls (F, G). *n* = 6 (D), *n* ≥ 13 (E, G), *n* = 5 (F). In (A, B, D, F), data are presented as mean ± SEM. Statistical significance was determined using a two-tailed unpaired *t* test (A–C, E, F) and two-tailed Mann-Whitney *U* test (D, G). Exact *P* values are indicated on the respective panels.

**Figure S7.**
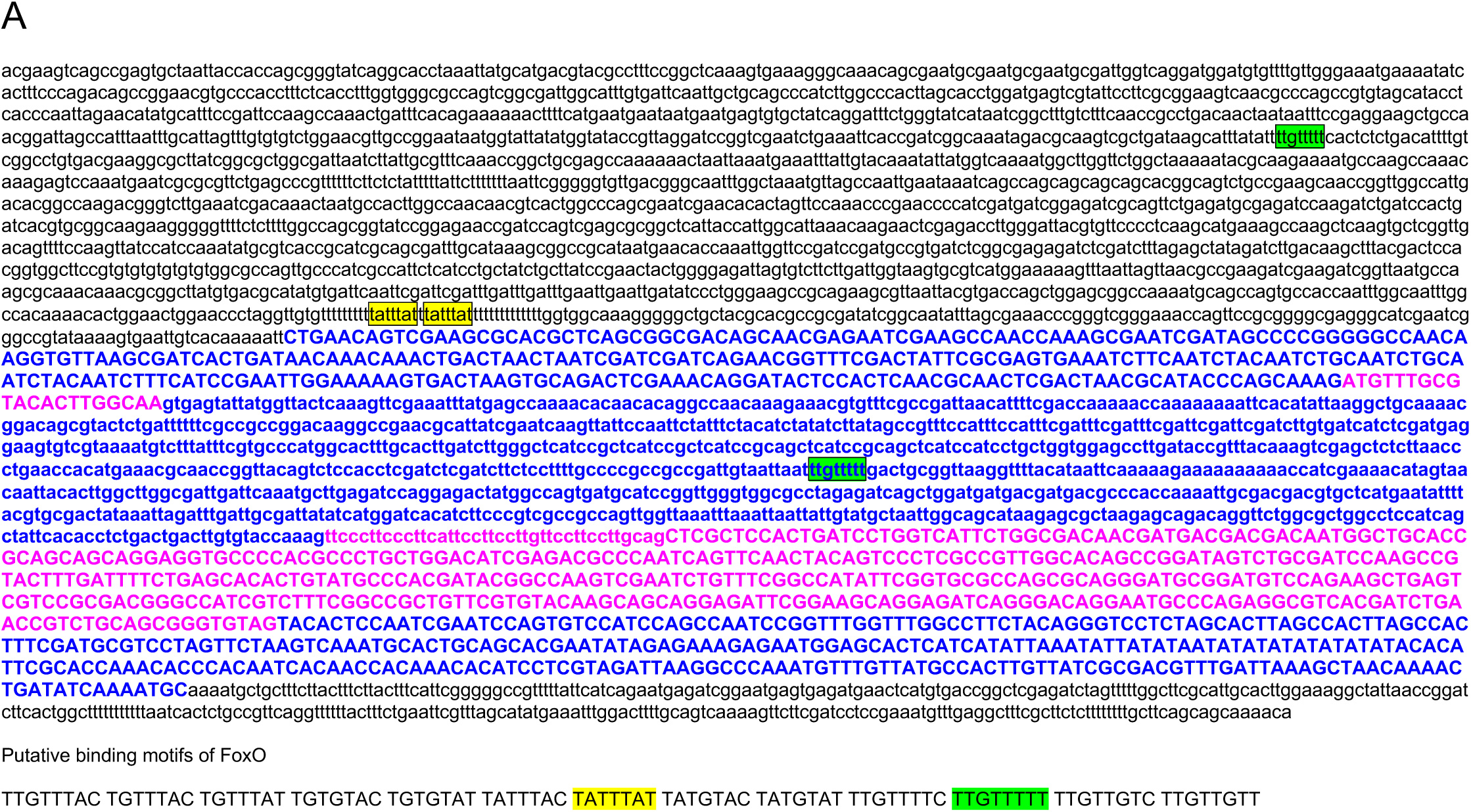
(related to. **Figure 7). Identification of putative FoxO-binding motifs within the *Lst* promoter region** (A) Schematic representation of putative FoxO-binding motifs within the 2 kb proximal promoter region of *Lst*, indicated by yellow and green boxes. For the sequence nomenclature: bold letters denote the *Lst* genomic sequence; uppercase letters represent exons; lowercase letters represent introns; pink text indicates the coding sequence (CDS); and lowercase letters in pink signify alternative splicing events.

**Figure S8.**
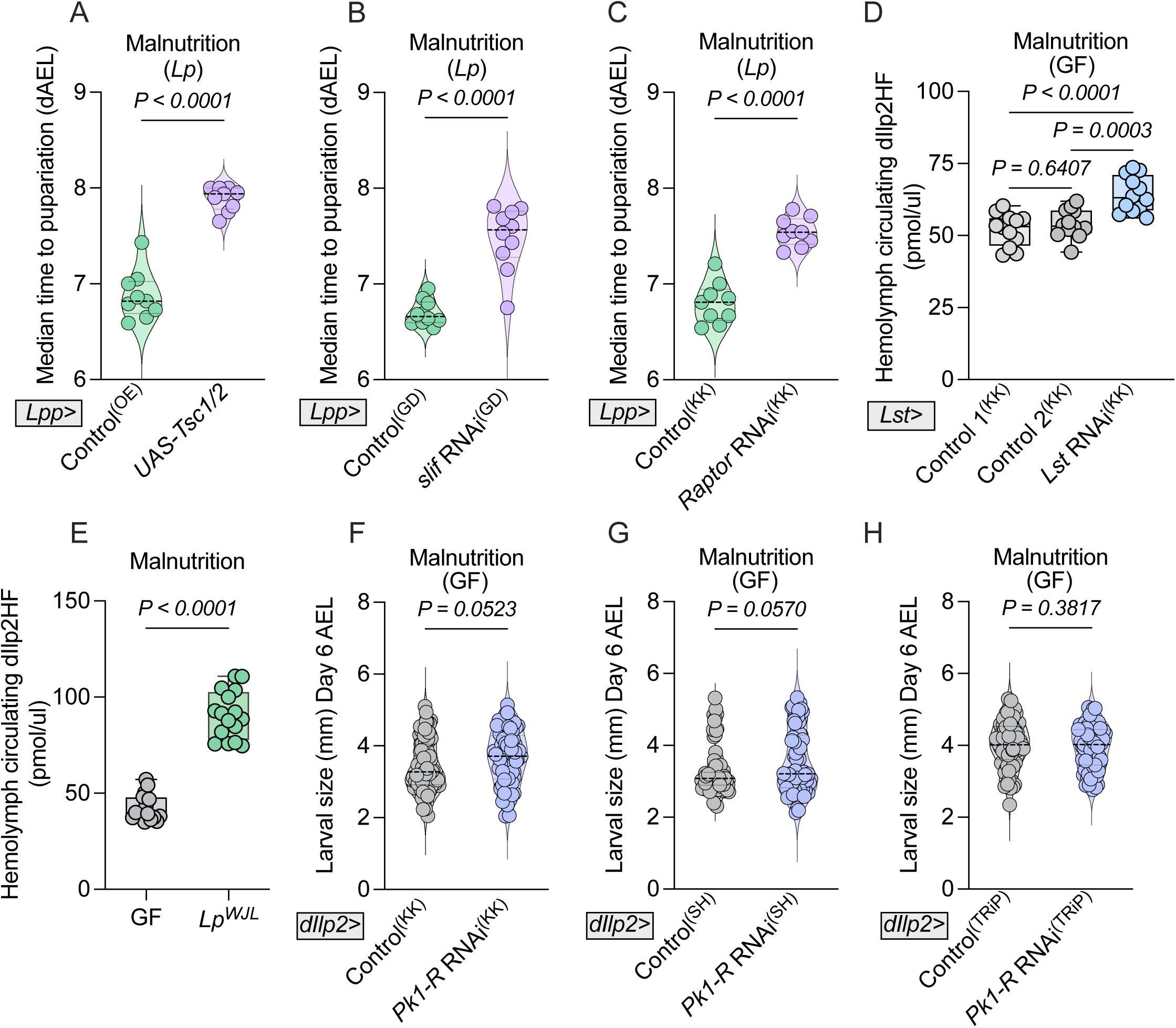
(related to Figure 8). Systemic TORC1 and Lst peptide coordinately regulate developmental timing and circulating insulin-like peptides. (A–C) Under malnutrition, the median time to pupariation was measured in *Lp*-associated larvae with fat body specific genetic manipulations, including: Overexpression of *Tsc1/2* (*Lpp>UAS-Tsc1/2*) compared to overexpression controls (A); knockdown of *slif* (*Lpp>UAS-slif* RNAi) compared to GD controls (B); or knockdown of *Raptor* (*Lpp>UAS-Raptor* RNAi) compared to KK controls (C). *n* ≥ 9. (D) Under malnutrition, circulating dIlp2(HF) in the hemolymph of germ-free larvae with *Lst* knockdown in *Lst*-expressing cells (*Lst>UAS-Lst* RNAi) and their respective KK controls were quantified by ELISA. *n* = 12. (E) The effect of *Lp-*association was assessed by measuring circulating dIlp2(HF) levels in the hemolymph of undernourished heterozygous *Ilp2¹* gd2HF larvae. *n* = 16. (F–H) Larval sizes were measured at Day 6 AEL in undernourished germ-free larvae reared with *Pk1-R* knockdown in IPCs (*dIlp2>UAS-Pk1-R* RNAi), compared to their respective controls. Results are shown for the KK RNAi line (F), the SH RNAi line (G), and the TRiP RNAi line (H). *n* ≥ 67. Statistical significance was determined using a two-tailed unpaired *t* test (A, C, H), a two-tailed Welch’s *t* test (B), a one-way ANOVA with Tukey’s multiple comparisons test (D), a Lognormal *t* test (E), or a two-tailed Mann-Whitney *U* test (F, G). Exact *P* values are indicated on the respective panels.

